# Scalable automated segmentation quantifies mitochondrial proteins and morphology at the nanoscale

**DOI:** 10.1101/2025.07.28.667278

**Authors:** Yuan Tian, Alexander Sauer, Mariia Dmitrieva, Hongqing Han, Hannahmariam T. Mekbib, Yujin Bao, Xiaojia Guo, Tian-min Chen, Robert Safirstein, Gary Desir, Jens Rittscher, Joerg Bewersdorf

## Abstract

Ultrastructural changes of mitochondria are closely associated with metabolic dysfunction, leading to a variety of human disorders. These changes can be visualized by pan-Expansion Microscopy (pan-ExM), a 3D microscopy technique requiring only standard fluorescence microscopes, at a throughput exceeding that of the current gold standard, 3D electron microscopy, by orders of magnitude. However, a lack of tools that enable the characterization and quantification of the observed ultrastructural features in the acquired 3D datasets at a comparable throughput has hindered the widespread adoption of pan-ExM for quantitative imaging. Here, we present an automated deep-learning based segmentation approach that utilizes pan-ExM’s power to acquire multi-channel images and uses specific labeling as the annotation for the training of the segmentation network. This ‘molecular annotation’ reduces the required manual annotation effort for mitochondria substructures to just a few hours when setting up the experiment and thereby provides access to 3D suborganellar morphology of mitochondria at an unprecedented throughput. Our approach, which we term MAPS (Mitochondrial Automated Pan-ExM Segmentation), enables for the first time to quantify mitochondrial ultrastructural morphology at scale. We demonstrate this power by characterizing the 3D mitochondrial morphology at the organelle and sub-organelle level in tens of HeLa cells under different treatments and localizing mitochondrial proteins in the sub-organellar context. To demonstrate our technology in tissue, we compare the ultrastructural morphology of mitochondria in proximal tubules of kidneys of mice exhibiting acute kidney injury (AKI) with those of untreated mice, revealing striking differences in their cristae structure. MAPS can easily be adapted to different cell and tissue types, allowing the analysis of tens of samples per day, and therefore provides a versatile tool for a comprehensive understanding of mitochondrial ultrastructural changes in many disease contexts. Requiring only standard fluorescence microscopes and computer infrastructure, MAPS is readily adoptable by any lab.

**Highlights:**

- MAPS utilizes specific labeling to train a segmentation model for 3D super-resolution pan-ExM images
- Suborganellar mitochondrial features and their changes in disease models are quantified at high throughput
- 3D protein distributions are correlated to ultrastructural features in mitochondria
- Requiring only standard lab infrastructure, MAPS is readily adoptable

## Introduction

Mitochondria play a critical role in cellular metabolism, stress responses, homeostasis maintenance and the regulation of cell death in eukaryotic cells^1–3^. Their specialized and multifaceted functions are linked to their ultrastructure, which exhibits heterogeneous morphology across different cell types and tissues^4^. Mitochondria are uniquely structured, with two proteolipid membranes that establish five distinct compartments: the outer mitochondrial membrane (OMM), intermembrane space (IMS), inner mitochondrial membrane (IMM), cristae, and matrix, arranged from the outermost to the innermost^5^. In contrast to the OMM which is relatively flat, the IMM is highly folded into cristae that provide a large surface area for the membrane-embedded ATP synthases. The organization of cristae controls the basic rate of cellular metabolism and is an indication of overall mitochondrial health^6,7^. Maintaining the mitochondrial ultrastructural integrity and organization is vital, as abnormal ultrastructural changes are strongly associated with functional deficiencies and the development of mitochondrial diseases. For example, disruptions and alterations in cristae organization are implicated in neurodegenerative, metabolic, and cardiovascular diseases^8–11^. Thus, understanding the relationships between mitochondrial structure and function in health and disease requires investigating mitochondrial molecular components and ultrastructural characteristics within their native cellular or tissue context.

Mitochondria feature a large range of different shapes and cristae morphologies, which are both constantly modified^8^. The observed shapes include compact, branching, elongated, and donut-shaped mitochondria, those featuring nanotunnels, and megamitochondria, as determined from 3D reconstructions using SBF-SEM^8^. Cristae are rapidly remodeled by key mitochondrial-shaping proteins, such as the mitochondrial contact site and cristae organizing system (MICOS) complex and dynamin-related protein optic atrophy protein 1 (OPA1)^11^. Other proteins also regulate the morphology and dynamics of mitochondria, orchestrating fusion and fission processes and participate in quality control^12–14^. Proteomic studies have identified over 1,100 mitochondrial proteins in human cells, with more than 40% linked to human diseases^15,16^. Mitochondrial diseases can manifest at any age, display any inheritance pattern, and involve any organ system, presenting with a wide range of clinical symptoms. The heterogeneity in the clinical manifestation of these diseases makes diagnosis and management extremely challenging^17^. High-resolution imaging tools that can quantify the distribution of proteins of interest in the context of mitochondria sub-organellar morphology at a reasonably high throughput will contribute to understanding these diseases.

Optical super-resolution (SR) microscopy technologies, including single-molecule localization microscopy (SMLM), stimulated emission depletion (STED) microscopy, has made it possible to systematically examine the 3D distribution of proteins at the sub-mitochondrial scale and investigate their specific functions^18–28^. These fluorescence-based technologies highlight, however, only the molecules that have been specifically labeled and struggle showing the ultrastructural context necessary to determine how specific proteins regulate mitochondrial shape and function. In contrast, volume electron microscopy (vEM) techniques facilitate the 3D visualization of mitochondrial ultrastructure but in general do not reveal the position of individual proteins therein. Correlative light and electron microscopy (CLEM) techniques overcome this dilemma by combining the global contrast of electron microscopy with the precise molecular localization provided by fluorescence microscopy, allowing to identify and map specific proteins in the context of suborganellar structure^29–31^. Recent advances, such as correlative cryo-SR/focused ion beam scanning electron microscopy (FIB-SEM), further improved ultrastructural preservation, imaging sensitivity and resolution^32^. However, the high complexity and cost of 3D CLEM techniques and their extremely low throughput severely limits their broad applicability.

Another optical SR technique, Expansion microscopy (ExM), has emerged over the last decade and demonstrated broad utility across diverse areas of biological research^33–46^. We recently developed pan-Expansion Microscopy (pan-ExM), an ExM variant that generates CLEM-like contrast using standard fluorescence microscopes^47,48^. pan-ExM expands hydrogel-embedded fixed cells or tissue 13 to 24-fold in each dimension, thereby improving the effective spatial resolution of standard diffraction-limited light microscopes (∼250 nm lateral resolution) down to ∼10 nm while preserving the sample’s protein content. The latter not only allows for the application of immunolabeling after expansion that will reveal the distribution of a specific protein population of interest, but also bulk (pan-) stainings of whole classes of molecules, e.g., using an NHS-ester conjugated dye that will covalently bind to primary amines present in essentially all proteins. The combination of a pan-staining with the ∼10 nm resolution achieved by the high expansion factor reveals, similar to the heavy-metal stain of classical EM, the general ultrastructural morphology of the sample and provides important context to the immunolabeled proteins. pan-ExM therefore addresses the limitations of CLEM by offering an easily accessible and time-efficient ultrastructure imaging method, especially when using high-end spinning disk confocal microscopy. For instance, it takes only about 15 minutes to acquire the complete volume of a HeLa cell in three colors (e.g. immunolabeling, nuclear staining and pan-staining) at a 3D resolution of ∼15×15×50 nm.

Our previous studies demonstrated that pan-ExM revealed the 3D ultrastructural features of mitochondria, showing that this technology is well-suited to investigate the distribution of mitochondrial proteins in context of mitochondria morphology. However, the large amount of data contained in each pan-ExM dataset poses extreme challenges for segmentation, classification and quantitative analysis of mitochondrial structure and protein distributions. Manual image segmentation and classification are labor-intensive, time-consuming and error-prone processes. Machine learning (ML)-based image analysis methods that have been increasingly applied to biological imaging in recent years, offer unbiased, reproducible, time-efficient and accurate automated processing of large, complex datasets^49^. Specifically, deep learning (DL)-based image analysis methods have been demonstrated for mitochondrial segmentation in both fluorescence microscopy^50–55^ and EM images^32,56–64^. These methods have been trained in a supervised way which bases their success on the availability of a large amount of annotated samples. However, the comprehensive annotation of 3D structures is inherently time-consuming and requires expert annotators to ensure accuracy^65^. The annotation requirement therefore poses a severe limitation on the application of DL techniques to new imaging technologies. The unique correlative imaging capabilities of pan-ExM that distinguish it from vEM, however, provide a remedy to the annotation challenge: images of specifically labeled structures acquired in one channel can be utilized during the training phase to automatically annotate the same structures in the simultaneously acquired pan-staining channel.

Here, we introduce “MAPS” (Mitochondrial Automated Pan-ExM Segmentation), a DL-based algorithm that automates the segmentation and classification of mitochondrial subcompartments from 3D pan-ExM images, enabling high-throughput quantification of morphological features and the distribution of specific proteins in their context. Taking advantage of the unique multichannel pan-ExM data, MAPS is initially trained using specific labeling of mitochondria as the annotation to automatically distinguish between mitochondria and the cytoplasm and other organelles. The resulting mitochondria-specific segmentation massively reduces the need for manual annotation. Human annotations, grounded in the knowledge of mitochondrial ultrastructure, are subsequently incorporated to refine the final model. After training, using this refined model, MAPS extracts mitochondrial subcompartments from 3D pan-staining images alone, eliminating the need for specific labels for their identification. After segmentation, MAPS allows users to quantify mitochondrial ultrastructural features. We demonstrate its capabilities by analyzing the effects of mitochondrial perturbations, revealing morphological changes in shapes and subcompartments. Furthermore, by utilizing pan-ExM’s compatibility with immuno-labeling mitochondria, MAPS facilitates the quantitative analysis of specific protein distributions in mitochondrial subcompartments. Revealing the changes in cristae organization in proximal tubules of mouse kidneys experiencing cisplatin (CP)-induced acute kidney injury, observed in >10,000 mitochondria extracted from 50 pan-ExM volumes, demonstrates automated ultrastructural mitochondrial analysis in disease research at a throughput and number dramatically exceeding previous vEM studies.

## Results

### MAPS workflow

The overall design and workflow of MAPS is schematized in **Figure 1**. Cell or tissue samples are processed using the pan-ExM protocol^47,48^ (**Figure 1A, STAR Methods**). Briefly, fixed samples, after incubating them in a solution of acrylamide monomers (AAm) and formaldehyde (FA), are embedded in a cleavable polyacrylamide hydrogel. The sample is then denatured and mechanically homogenized using sodium dodecyl sulfate (SDS) and heat. After expanding it 4 to 5-fold in deionized water, the gel is additionally embedded in a neutral hydrogel that acts as a scaffold to maintain its expanded state, and then in a third non-hydrolyzable hydrogel to allow for a later second round of expansion. Next, the crosslinkers of the first and second hydrogels are cleaved using sodium hydroxide and the hydrogel is neutralized with PBS. Specific mitochondrial proteins can then optionally be labeled using antibodies or dyes while the application of a pan-staining provides structural context. The mitochondrial matrix, in particular, is stained strongly by NHS-ester dyes. Finally, the hydrogel is expanded to its final size in deionized water, resulting in a linear expansion factor of 14 to 18. These samples are then imaged in 3D using a standard confocal or spinning disk confocal fluorescence microscope.

**Figure 1.**
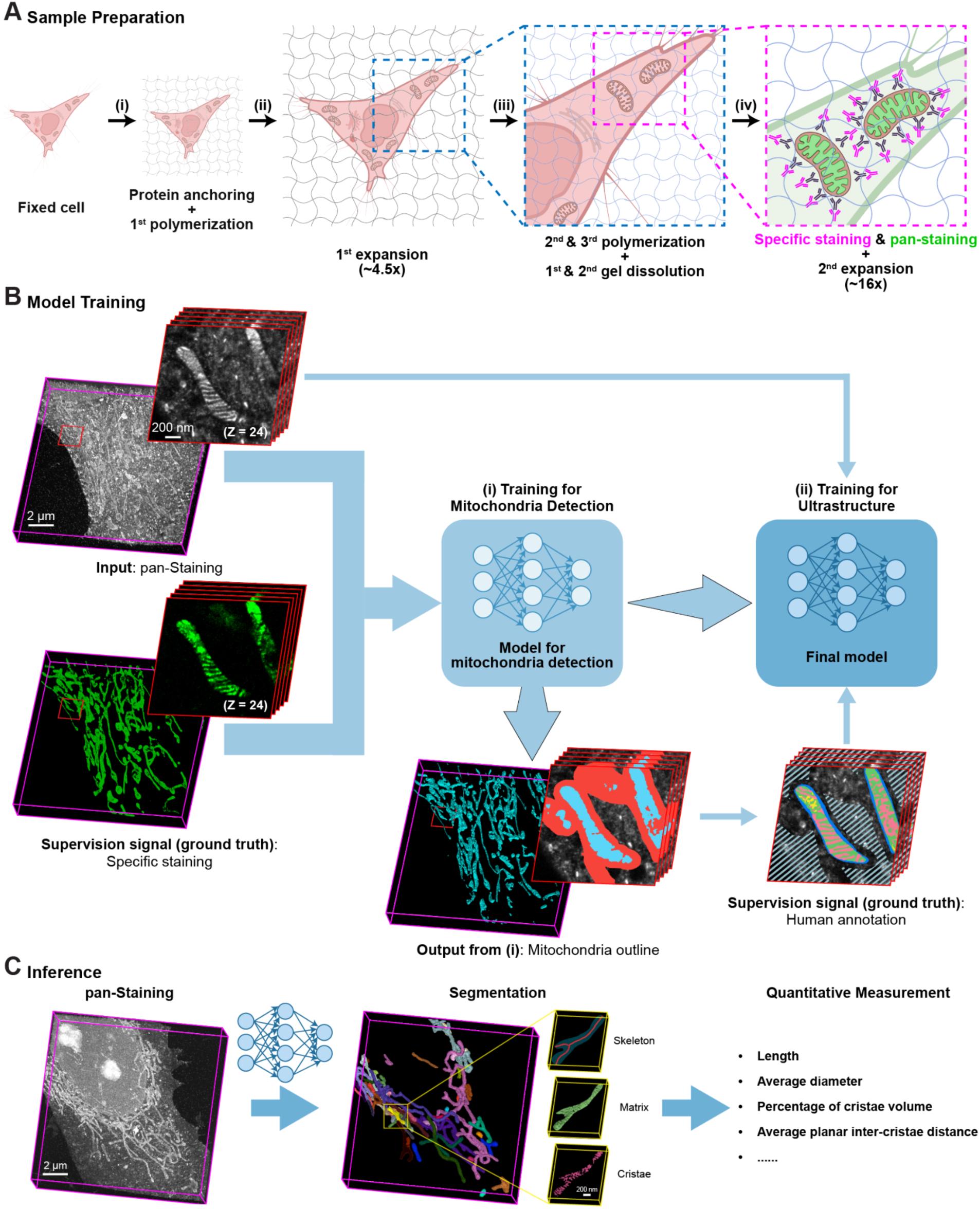
MAPS workflow. (A) Schematic overview of the pan-ExM protocol achieving ∼16-fold iterative expansion and allowing it to visualize specifically labeled structures or proteins within the ultrastructural context of the cell. Clipart created with BioRender.com. (B) Computational pipeline for model training. (i) Specific labeling of the mitochondrial ultrastructure (e.g., MitoTracker shown in green) serves as the supervision signal (“ground truth”) for training a segmentation model that predicts the 3D outline of mitochondria (red) and their matrix (cyan) from 3D pan-stained data sets (gray). (ii) The model is further trained for ultrastructural detection by human annotations of a small subset of the pan-stained training data including 25 cross-sections of mitochondria (e.g. the magnified region shown in the red box). This refinement enables the prediction of mitochondrial substructures. Scale bars have been corrected for the expansion factor. (C) The final model resulting from the training facilitates automated large-scale segmentation and quantitative analysis of mitochondria based solely on the pan-staining image. Scale bars have been corrected for the expansion factor.

To enable our DL network to segment mitochondria subcompartments from 3D data sets of the pan-staining channel alone, we train a model in a two-step process in which we take advantage of two-color image data where the pan-staining channel is augmented by a second channel showing mitochondria-specific fluorescence labeling (**Figure 1B**).

The first training step utilizes this second channel to train an initial model to segment the mitochondria volume from the pan-staining channel. This step does not require any human annotations and, thanks to the absence of this bottleneck, can take advantage of large numbers of 3D data sets that can be quickly acquired by the confocal microscope (e.g. 50 two-color data sets of 2048×2048×200 voxels acquired in 3 hours). If the signal shown in the second channel covers the target structure densely and features low off-target labeling (e.g. MitoTracker), simple denoising methods and traditional image-processing operations (smoothing, dilation) as described in the **Methods Section** allow the algorithm to derive target segmentation masks with sufficient accuracy. These target masks are subsequently used for supervised training. In the case of immunolabeling which shows substantial off-target background signal and sparser mitochondrial labeling, the problem is cast into a semi-supervised learning problem by ignoring pixels in close neighborhoods of the antibody localizations. The details are described in the **Methods Section**.

In the second step, we refine the model training by building on the mitochondria outlines obtained in the first step and filling in ultrastructural details based on manual annotations. For this purpose, the mitochondrial volume is divided into the subcompartments of ‘intermembrane space’ (IMS, blue), ‘matrix’ (green), and ‘cristae’ (pink). The IMS volume is defined as the space between the OMM and the inner boundary membrane (IBM), while the cristae volume is defined as the intercristae space (**Figure 1Bii**). This fine-tuning step works very efficiently as the segmentation task is substantially simplified because the model can focus exclusively on the ultrastructure as all background structures within the cell are masked out by the outline model. Moreover, the model training in the first step results in a data representation that only requires selected annotation of the data to fine tune the model (see **Figure S1**).

The presented pipeline is in general agnostic to the network architecture of the model but we find the best results provided by a well-calibrated 3D convolutional UNet^66,67^. Moreover, the UNet architecture allows to naturally mirror the two-step training pipeline by enabling the reuse of the feature-extracting encoder from the first step for the ultrastructure segmentation and only add a fine-tuned decoding branch. This approach therefore allows us to substantially decrease the required total annotation time to less than a day (**Figure S1**).

In order to validate the segmentation quality, the segmentation masks of 30 identified mitochondria instances in expanded HeLa cells were proof-read and corrected by a human annotator. Comparing the original predictions with the corrected versions results in an intersection-over-union (IoU) score of 0.97 which confirms the high reliability of the model.

The final model can subsequently be applied to segment mitochondria and their ultrastructure from pan-staining data sets alone (**Figure 1C**). The resulting segmentation masks then allow for the extraction of metrics of interest, such as the volume fraction occupied by the cristae, and form the base to skeletonize the mitochondria. The resulting skeletons reveal the topology of the mitochondria and enable additional measurements including the length and diameter of the mitochondria.

### MAPS enables quantitative analysis of mitochondrial morphology in HeLa cells

To validate the performance of MAPS, we first applied it to quantify mitochondrial shape and ultrastructure in 3D pan-ExM images of expanded HeLa cells. Using the pan-staining channel as the sole input, MAPS successfully identified and differentiated mitochondria from the cytosol background, correctly identifying 94% ± 2.6% of the mitochondrial volume (mean ± SD; n=15 imaged volumes; **Figure S2, STAR Methods**), and further classified them into distinct compartments: the volume of IMS (blue), matrix (green), and cristae (pink). **Figure 2A** shows a volumetric view of pan-stained mitochondria extracted from a 3D stack image in an expanded HeLa cell (**Video S1**). After correcting for the expansion factor of the experiments in HeLa cells (**Figure S3A**, mean ± SD: 14.8 ± 0.5), the achievable lateral and axial resolution using a high-end confocal microscope with a 60X/1.2NA water immersion objective is ∼17 nm and ∼70 nm, respectively. During segmentation, areas where the 3D distance between neighboring cristae is below this resolution limit were challenging to classify as either cristae or matrix. These regions, along with spaces presumably occupied by mitochondrial nucleoids^68,69^ (**Figure S4**), were designated as ambiguous volumes in yellow. Notably, for 98% of analyzed mitochondria this ambiguous volume amounted to less than 5% (**Figure S3B**), indicating that the ambiguous classification had minimal impact on the overall analysis.

**Figure 2.**
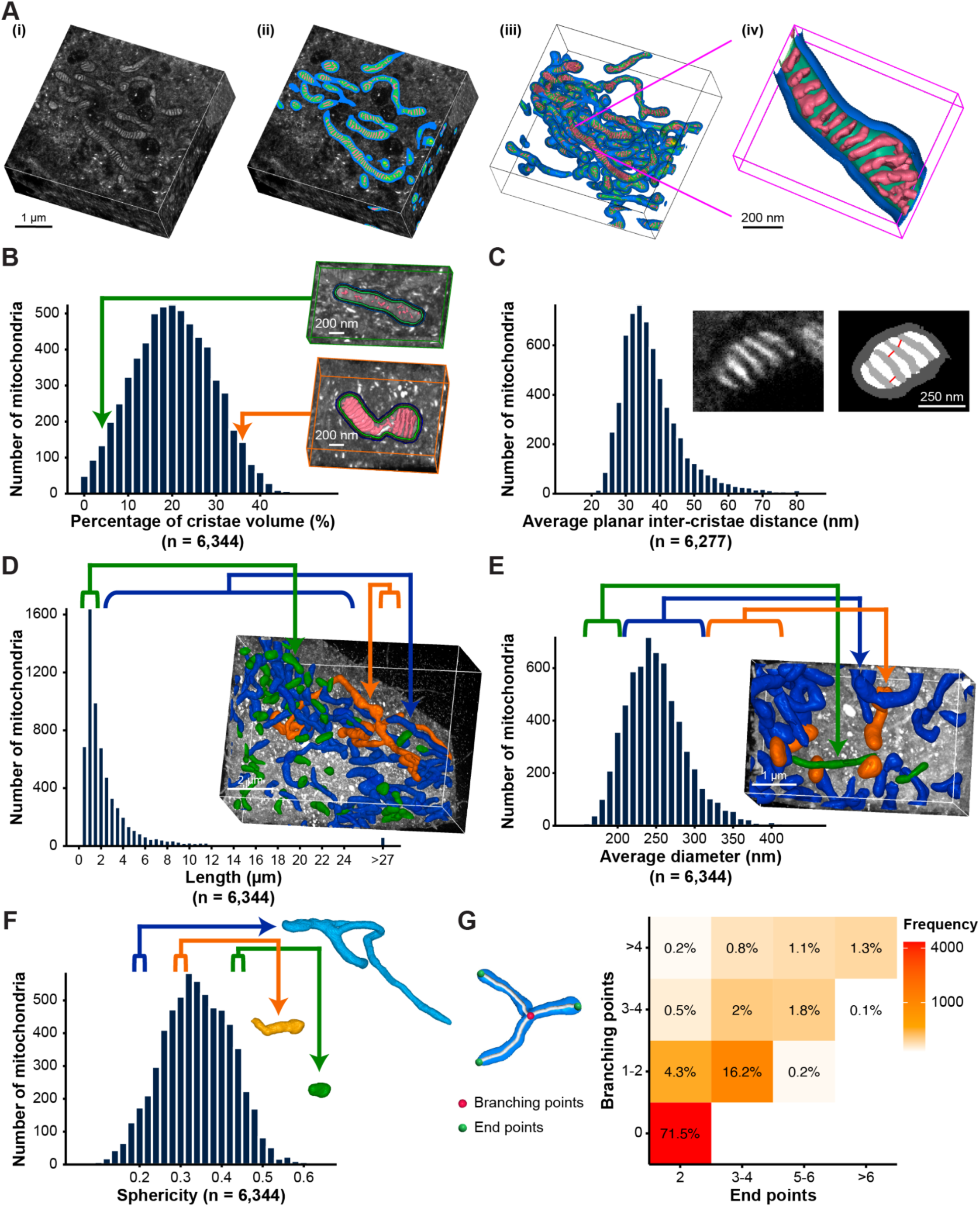
Quantitative analysis of mitochondrial structure and ultrastructure in HeLa cells. **(A)** Representative segmentation results generated by MAPS: (i) 3D pan-ExM dataset of an expanded HeLa cell. (ii) Segmentation of mitochondrial substructures, including the intermembrane space (IMS, blue), matrix (green), cristae (pink), and ambiguous volume (yellow). (iii) Volumetric view of the reconstructed mitochondria. (iv) Magnified view of a selected region in magenta. Scale bars have been corrected for the expansion factor. **(B-F)** Quantitative analyses of mitochondrial morphology (*n* = 6,344 mitochondria, *N* = 30 cells from 4 independent experiments): (B) Distribution of cristae volumes as a fraction of the volume enclosed by the inner boundary membrane (IBM) (mean ± SD: 20.3% ± 9.0%). Insets display mitochondria with cristae fractional volumes of 5% (green frame) and 36% (orange frame), with cristae highlighted in pink. (C) Average planar inter-cristae distance (mean ± SD: 39 ± 9.8 nm); insets show raw data from a single mitochondrion (left) and its segmentation (right), with red lines indicating planar inter-cristae distances. (D) Mitochondrial length (*l*) distribution (mean = 3.3 µm). The inset shows a 3D dataset where mitochondria with *l* < 1.5 µm are shown in green, those with *l* > 27 µm are orange, and those in between are blue. (E) Average mitochondrial diameter (*d*) (mean ± SD: 250 ± 40 nm). The inset shows mitochondria with *d* < 200 nm shown in green, those with *d* > 300 nm in orange, and those in between in blue. (F) Mitochondrial sphericity (mean ± SD: 0.34 ± 0.09). Insets illustrate mitochondria with sphericity values of 0.19 (blue), 0.31 (orange) and 0.45 (green). Measurements and scale bars were corrected for the expansion factor. **(G)** Heatmap illustrating the correlation between the number of mitochondrial branching points (red spheres) and end points (green spheres) (*n* = 6,344 mitochondria, *N* = 30 cells from 4 independent experiments).

To assess cristae density, we quantified the percentage of cristae volume from 6,344 mitochondria, revealing an average volume proportion of 20.3% with a large mitochondria-to-mitochondria variation (SD: 9.0%) (**Figure 2B**). The cristae-to-cristae planar distance in HeLa cells was measured by calculating the shortest planar distance from one cristae invagination to its closest neighbor, resulting in an average value of 39 ± 9.8 nm (mean ± SD) from 6,277 mitochondria (n = 30 cells), with only 10% exceeding 50 nm (**Figure 2C**).

The segmentation data obtained from MAPS enabled us to extract several key quantitative measures of mitochondrial shape (length, diameter and volume) and morphological complexity (sphericity, branching network). Mitochondrial length was measured using skeleton analysis from 6,344 mitochondria in 30 HeLa cells, showing a broad, asymmetric distribution with an average value of 3.3 µm and a median value 1.7 µm with the middle 80% of mitochondria in the distribution ranging from 0.7 µm to 10.4 µm in length (**Figure 2D**), which is close to previously reported FIB-SEM data from two HeLa cells (mean ± SD: 2.3 ± 3.2 µm)^62^. For the same mitochondria, we calculated their diameters as twice the mean distance from the outer membrane to the corresponding skeleton. The obtained diameters feature a nearly symmetric distribution with an average value of 250 ± 40 nm (mean ± SD) from 6,344 mitochondria (n = 30 cells) (**Figure 2E**). This value is on the lower end but in agreement with reported values of 200 and 700 nm in a range of mammalian cell types^70^. Moreover, we observed a positive correlation between the cristae volume fraction and mitochondrial diameter (p < 0.001; **Figure S3D**), suggesting that thicker mitochondria generally contain more cristae. In contrast, no significant correlation is observed between mitochondrial length and cristae volume fraction (p = 0.49; **Figure S3E**). Mitochondrial volume was measured by calculating the overall volume of the segmentation mask of each segmented mitochondrion (**Figure S3F**). Analogous to the length measurements, the volume distribution exhibits a prominent tail with an average value of 0.13 µm^3^, a median value of 0.07 µm^3^, and 80% of mitochondria lying within 0.03 µm^3^ and 0.25 µm^3^. These values for the mitochondrial average volume are about half of those previously reported for much smaller sample sizes (mean ± SD: 0.29 ± 0.54 µm^3^; 421 mitochondria across two HeLa cells) in FIB-SEM datasets^62,71^. A possible reason for this discrepancy could be that the two cells imaged in the FIB-SEM experiment were not representative of the cell population which could have been heterogeneous. The mitochondria in our data sets regularly formed large networks and on average only 6 ± 4.5 (mean ± SD) mitochondria contained 20% of the overall mitochondria volume per cell even though the average number of mitochondria per cell is found to be 211 ± 80 (mean ± SD), which captures how much of the total mitochondria volume per volume of view is concentrated in only a few mitochondria, reflects this imbalance with its value of 0.53 ± 0.09 (mean ± SD; value approaching 1 corresponds to the presence of dominating networks) (**Figure S3G**). Additionally, to measure the overall mitochondrial volume present in an imaged volume of view (VOV), we calculated the fraction of the cytoplasm occupied by mitochondria, resulting in a value of 9% ± 2% (30 VOVs; **Figure S3C**). Mitochondrial sphericity varies with cellular stress, metabolic conditions and the cell cycle^4,14,72–78^. The sphericity is derived from the volume and surface area^79^ resulting in an average value of 0.34 ± 0.09 (mean ± SD) from 6,344 mitochondria (n = 30 cells) with 1 representing a perfect sphere (**Figure 2F**). To assess the mitochondrial topology, we extracted branching points and end points from the skeletons. In line with the previously reported metrics, most mitochondria (71.5%) show a simple shape with two end points and no branching points and only about 4% show complex topologies with more than 5 end points (**Figure 2G and S3H**). Additionally, a positive correlation was observed between the number of branching points and mitochondrial length, suggesting that longer mitochondria typically exhibit more branching (p < 0.001; **Figure S3I**).

### MAPS can localize proteins in mitochondrial subcompartments

Understanding mitochondrial biogenesis and function requires not only morphological analyses but also determining both protein composition and localization within mitochondrial subcompartments. To demonstrate MAPS’ capability to extract this information, HeLa cells were immunolabeled with three antibodies, each specific to different mitochondrial subcompartments. TOM20, a receptor protein in the translocase of the outer mitochondrial membrane (TOM) complex, is embedded in the OMM and facilitates the binding and import of nuclear-encoded proteins into mitochondria^80–83^ (**Figure 3A**). MIC60, the largest subunit of the mitochondrial contact site and cristae organizing system (MICOS) complex, is enriched at crista junctions, where it interacts with other subunits to control the formation of cristae junctions (CJs) and link them to the OMM through the formation of membrane contact sites^68,84–89^ (**Figure 3B**). COX4, the nuclear-encoded subunit of the cytochrome c oxidase (COX) complex, is located in the inner mitochondrial membrane and plays a key role in the assembly and stabilization of COX, as well as in cytochrome c docking^90–92^ (**Figure 3C**). Using MAPS, we identified antibody-labeled signal clusters for each protein, correlating them with segmented mitochondrial subcompartments. We first quantified the percentage of signal clusters within each 3D subcompartment volume, disregarding cytoplasmic background (**Figure 3D**). In agreement with TOM20 being a protein of the OMM, it was clearly enriched (84%) in the narrow IMS compartment between IBM and OMM (including OMM), with the remaining 16% attributed to non-specific antibody binding inside the IBM. MIC60, which is expected to localize at CJs, which for our compartmentalization should be located at the interface between the IMS and the matrix volume, was found for 95% of clusters in these two volumes. COX4, as an IMM marker, is expected to localize at the interface between cristae and matrix volume, and 68% of clusters were distributed between these two volumes. As an alternative analysis, we measured the relative position of the antibody signal to the IBM (**Figure 3E**). As expected, the majority of the TOM20 signal is localized clearly on the cytoplasmic side of the IBM, while MIC60 is centered on the IBM, and COX4 is located on the inside of the IBM. Additionally, we identified putative cristae junctions as the interfaces between the IMS and cristae segmentation volumes and measured the distance to the closest labeling cluster (**Figure S5**). In agreement with MIC60’s reported localization at cristae junctions, we find it significantly (p < 0.001) closer to the identified interfaces than TOM20 (**Figure 3F**).

**Figure 3.**
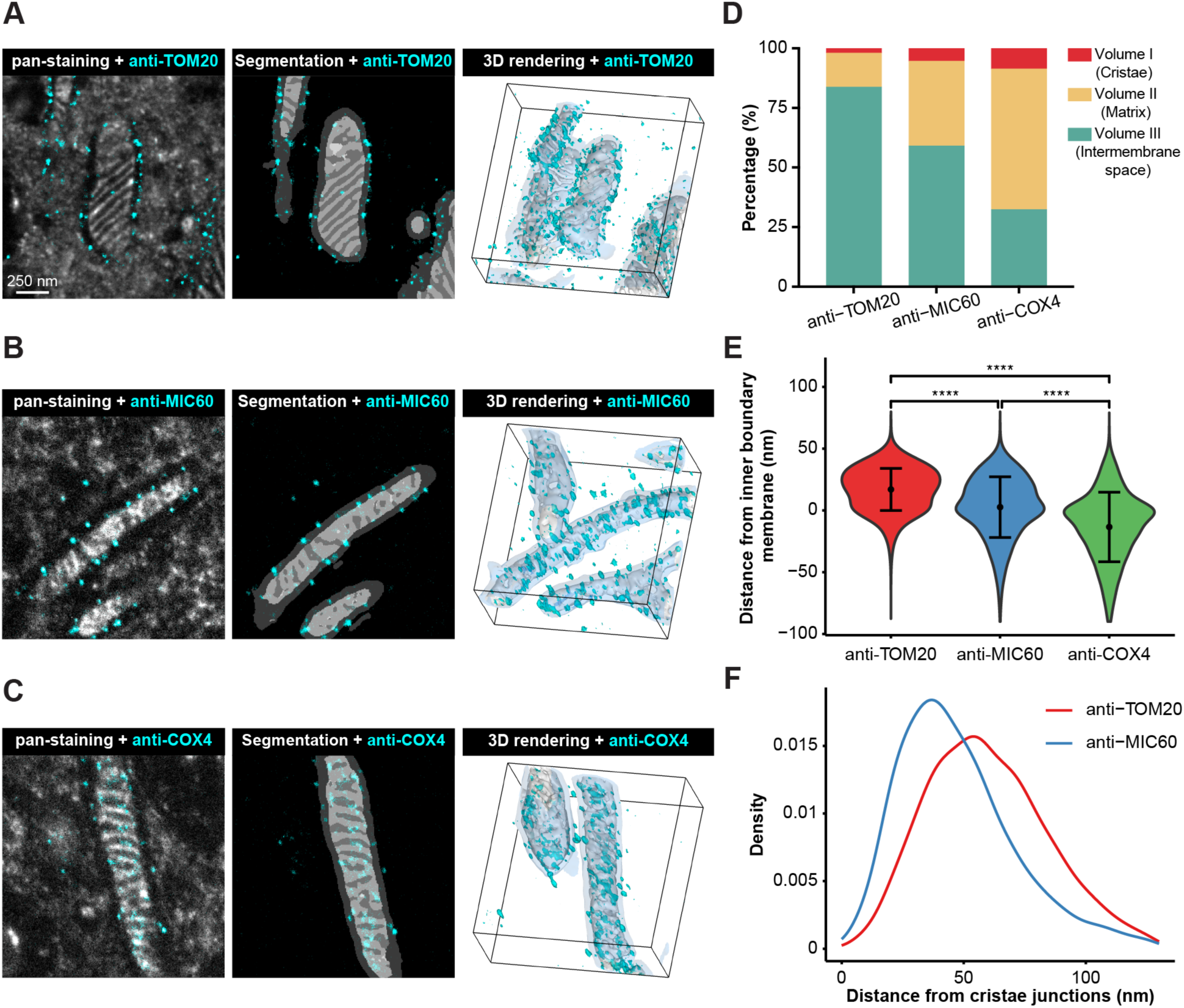
Quantitative analysis of protein localization in mitochondrial subcompartments. **(A-C)** Representative images of HeLa cells immunolabeled for mitochondrial proteins: TOM20, localized to the mitochondrial outer membrane (A); MIC60, a component of the mitochondrial contact site and cristae organizing system (MICOS) complex localized to cristae junctions (B); and COX4, localized to the cristae membrane (C). Immunolabeling data for these proteins (cyan) are overlaid with the corresponding raw pan-staining images (gray; left), segmentation results (gray; middle), and 3D-rendered volumes (gray; right), confirming the expected localization. Scale bar has been corrected for the expansion factor. **(D)** Quantitative analysis of antibody distributions of TOM20 (32,604 labels from 833 mitochondria in 3 HeLa cells), MIC60 (22,729 labels from 320 mitochondria in 3 HeLa cells), and COX4 (15,007 labels from 193 mitochondria in 2 HeLa cells) in 3D individual mitochondria. Results show that 84% of TOM20 localizes within volume III (IMS, which includes the outer membrane), 95% of MIC60, which is expected to localize at the interface between volumes II (matrix) and III, is distributed between these two volumes, and 68% of COX4, which is expected to localize at the interface between volumes I (cristae) and II, is distributed between these two volumes. **(E)** Violin plots showing the distance of antibody labeling from the inner boundary membrane (IBM). Negative values indicate signal localization on the matrix/cristae side of the interface. MIC60 is centered to the inner membrane, while COX4 localizes mostly inside and TOM20 outside of it. ****, P < 0.0001, Mann-Whitney Test. Measurements were corrected for the expansion factor. **(F)** Distribution of distances between antibody labeling and cristae junctions (identified as the interface between volumes I and III), showing that MIC60 localizes significantly closer to identified interfaces than TOM20 (P < 0.0001, Mann-Whitney Test). Measurements were corrected for the expansion factor.

### MAPS allows for detecting ultrastructural changes of mitochondria in perturbed cells

Mitochondria undergo profound morphological changes in response to metabolic activity. To evaluate the versatility and robustness of MAPS in identifying various mitochondrial states, we analyzed the changes in mitochondria morphology in HeLa cells treated with oligomycin, a well-known ATP synthase inhibitor. This treatment has been reported to modulate mitochondrial ATP levels, the pH in the matrix, membrane potential, and triggers mitochondrial fission^93,94^.

MAPS successfully identified and segmented mitochondria in both control and treated cells (**Figure 4A-B; Video S2 and S3)**, enabling quantitative comparison at the subcompartment level. Analysis of 8,067 mitochondria from 46 control cells and 15,329 mitochondria from 50 oligomycin-treated cells revealed substantial morphological changes. Oligomycin-treated mitochondria were significantly (p < 0.0001) shorter **(Figure 4E)**, but not significantly thicker or thinner (p = 0.76; **Figure 4G**), resulting in overall rounder, less voluminous mitochondria, compared to the control **(Figure 4F and Figure S6A)**. In contrast to the observed changes at the level of individual mitochondria, the total volume occupied by mitochondria within the cells did not change significantly (**Figure S6B**). Taking advantage of the obtained compartmental segmentation, we quantified changes at the level of cristae: the volume fraction of cristae per mitochondrion decreased significantly (p < 0.0001) in treated cells from 21% to 16% (**Figure 4C**). This decrease was accompanied by a simultaneous increase in the average planar inter-cristae distance from 40 to 46 nm (**Figure 4D**) and in the volume fraction of the mitochondria volume assigned as ambiguous by MAPS from 1.6% to 5.4% (**Figure S6C**). We interpret the latter as a partial shift of cristae morphology from a sheet-like to a more tubular appearance, which is more difficult for MAPS to segment.

**Figure 4.**
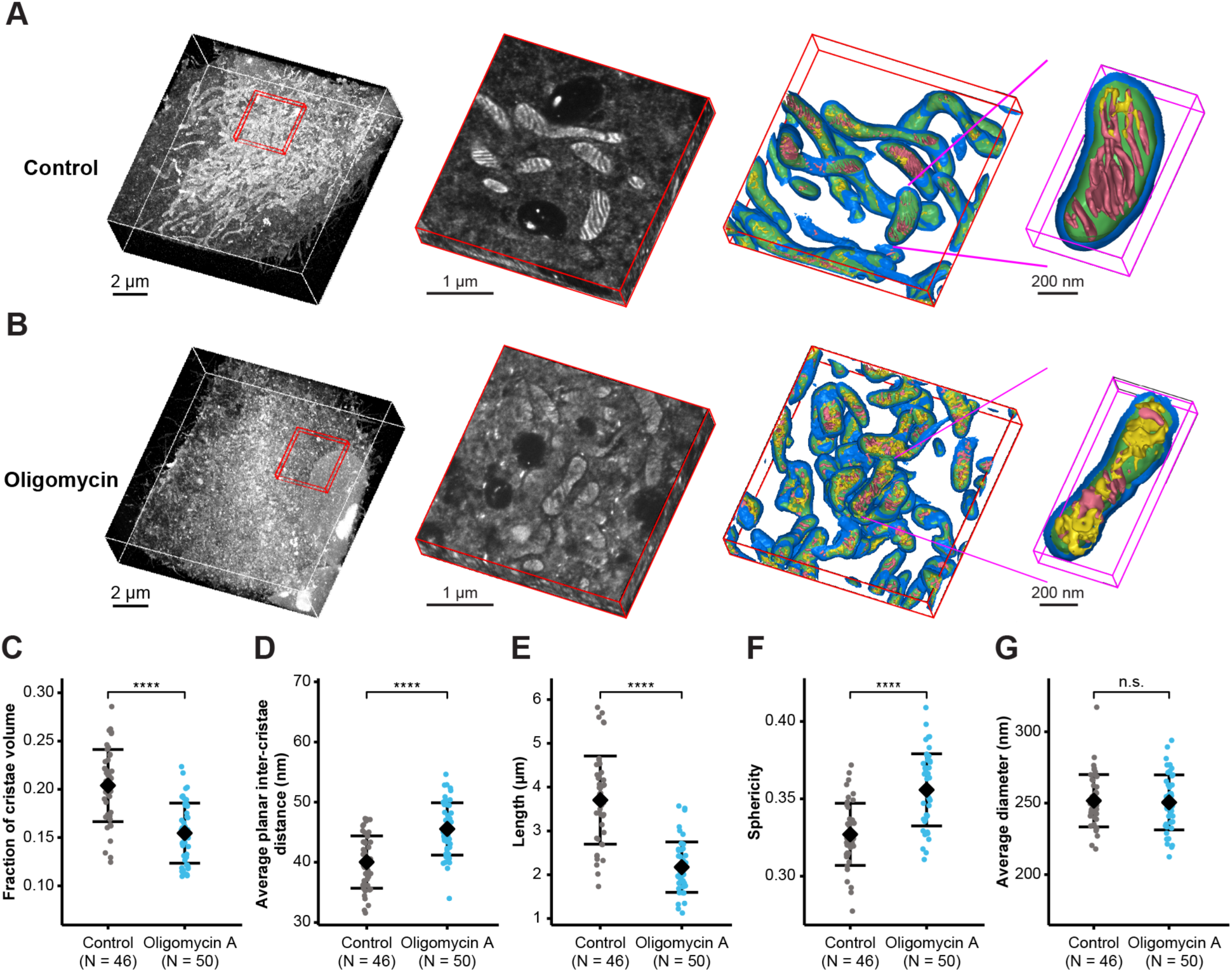
MAPS reveals organellar and suborganellar morphological changes in mitochondria in oligomycin A-treated HeLa cells. **(A and B)** Representative pan-ExM 3D image data and MAPS results of HeLa cells under different conditions: control (A), treatment with 10 μM oligomycin A (B). For each condition, the panels from left to right show: a volumetric view of an expanded HeLa cell; a magnified region of the cell (red box); a volumetric reconstruction of mitochondrial ultrastructure within the magnified region (blue: IMS; green: matrix; pink: cristae; yellow: ambiguous volume); and a magnified view of an individual mitochondrion. Scale bars have been corrected for the expansion factor. **(C-G)** Quantitative analysis of mitochondrial morphological changes under the different conditions (control: *n* = 8,067 mitochondria, *N* = 46 cells; oligomycin A: *n* = 15,329 mitochondria, *N* = 50 cells. Each data point represents the average from one cell. All derived from 6 independent experiments). Estimated means are shown in black, with error bars representing one standard deviation. ****, P < 0.0001 (Mann-Whitney Test); n.s., not significant. Oligomycin A treatments result in significantly less visible cristae, a larger inter-cristae distance, and significantly shorter and rounder mitochondria. Measurements were corrected for the expansion factor.

Our findings are consistent with previous studies, which reported fragmented mitochondrial networks, rounded mitochondria, and reduced cristae density following oligomycin treatment^93,95–97^. Additionally, we observed a significant (p < 0.0001) reduction in the Gini coefficient, indicating a more even distribution of volume between mitochondria and fewer large networks in oligomycin-treated cells (**Figure S6D**). Oligomycin treatment also resulted in fewer branching and end points per mitochondrion, along with a reduction in the length per branching point (**Figure S6E-G**).

### Quantitative analysis of mitochondrial morphology in mouse kidney sections

To demonstrate the broader application of MAPS, we applied it to mouse kidney tissue samples. The kidney, in particular its proximal tubules, contain large amounts of mitochondria that are essential for kidney function due to the high energy demand in electrolyte reabsorption^98,99^. Their dysfunction is indicative of renal disease and injury^100,101^. Proximal renal tubules are divided into proximal convoluted and proximal straight tubules, classified as S1, S2 and S3 segments based on subtle differences in cellular ultrastructure and function^102–104^. Segment S1/S2 of the proximal tubules, which we focus on in this study, contains a higher density of mitochondria. This aligns with their significant energy need for active solute transport, reabsorption, and related metabolic activities^105–109^.

We adapted a protocol for pan-ExM of tissue (pan-ExM-t) that we previously had developed for mouse brain sections^40^ for ∼50-µm thick vibratome sections of mouse kidney (see **STAR Methods**). The modified protocol resulted in ∼18-fold linearly expanded sections showing negligible distortion (**Figure S7A**, **S7B** and **S8**) and enabled pan-staining of mitochondria and immunolabeling of specific mitochondrial proteins (**Figure S7C**) in kidney tissue.

Not surprisingly, due to the different underlying cellular morphology, our model trained on HeLa cells alone did not reliably identify mitochondria in these kidney data sets. We therefore repeated the first training step designed to identify mitochondria. Since the application of live-cell dyes such as Mitotracker is difficult in tissue, we used mitochondria-specific immunolabeling as the annotation channel (**Figure S9A, STAR Methods**). As in the HeLa cell case, this did not require any manual annotation. Since mitochondrial subcompartments are essentially conserved across tissue types and model organisms, we then used the final model trained on HeLa cells to complement the newly trained model and refine the training to identify mitochondrial subcompartments in kidney samples (**Figure S9A**). This combination of old and new models allowed us to minimize manual proof-reading and corrections. An afternoon’s effort of a single human annotator was sufficient to yield satisfying results on automatically segmenting mitochondria and their subcompartments in wild-type mouse kidney sections using this retrained network (**Figure S9B-C, STAR Methods**).

**Figure 5A** shows an example of a single optical section from a 3D data set of a cortical proximal tubule. MAPS successfully automated the segmentation and classification of mitochondrial compartments in the 3D pan-ExM-t images (**Figures 5B** and **S8D; Video S4**). Quantification of mitochondrial length, diameter, and volume was performed using the same approach as described above. Measurements from 14,065 mitochondria across 32 expanded cortical proximal tubules revealed a broad, asymmetric distribution in mitochondrial length, with an average value of 2.6 µm and a median value 1.7 µm with the middle 80% of mitochondria in the distribution ranging from 0.9 µm to 5.0 µm. The average diameter was 436 ± 83 nm, and the volume was 0.29 ± 0.27 µm^3^ (mean ± SD) (**Figures 5C-E, S10A**). While the length is in rough agreement with that obtained from our HeLa cell data, mitochondria are nearly twice as thick, explaining the more than twice larger average volume per mitochondria. This is also reflected by a larger sphericity value of 0.39 ± 0.08 (mean ± SD), indicating that kidney mitochondria are slightly rounder than those in HeLa cells (**Figure S10B**). Further submitochondrial analysis showed that cristae accounted for 19.9% ± 9.7% (mean ± SD) of the volume enclosed by the IBM (**Figures 5F** and **G**), nearly identical to the value obtained in HeLa cells. The average planar distance between cristae was measured at 31.4 ± 8.0 nm (mean ± SD) (**Figure 5H**), about 20% smaller than in HeLa cells.

**Figure 5.**
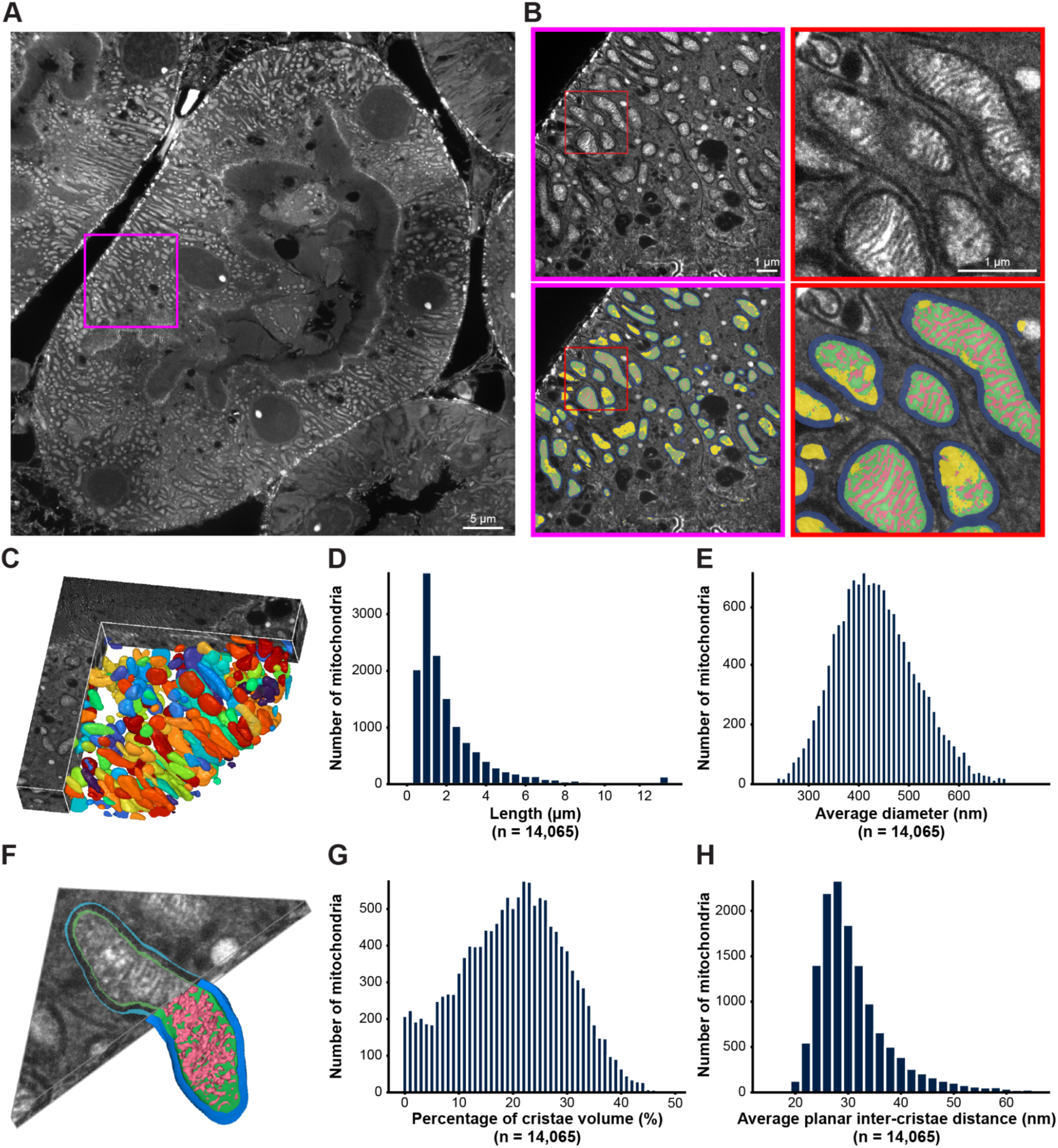
MAPS enables the quantitative analysis of 3D mitochondrial morphology in mouse kidney tissue. **(A and B)** pan-ExM enables visualization of mitochondria and their ultrastructure in ∼18-fold expanded mouse kidney sections. (A) Representative 3D image of an expanded proximal renal tubule. (B) Top left, a magnified view of the region highlighted by the magenta box in (A), with a further zoomed-in view of the area in the red box (top right). Bottom, the same regions with segmentation overlay, showing the IMS (blue), matrix (green), cristae (pink), and ambiguous volume (yellow). **(C)** Volumetric view of mitochondria in 3D pan-ExM images of a mouse kidney section. Scale bars have been corrected for the expansion factor. **(D and E)** Histograms displaying the distributions of mitochondrial lengths (mean = 2.6 µm) and average diameters (mean ± SD: 436 ± 83 nm). A total of *n* = 14,065 mitochondria were analyzed from 47 imaged volumes, and *N* = 32 proximal renal tubules are sampled from 6 independent experiments. Measurements were corrected for the expansion factor. **(F)** Volumetric view of the reconstructed mitochondrial ultrastructure (colors as in B). **(G and H)** Quantitative analysis of cristae volumes as a fraction of total mitochondrial volumes (mean ± SD: 19.9% ± 9.7%) and average planar inter-cristae distance (mean ± SD: 31.4 ± 8.0 nm). A total of *n* = 14,065 mitochondria were analyzed from 47 imaged volumes; *N* = 32 proximal renal tubules sampled from 6 independent experiments. Measurements were corrected for the expansion factor.

### MAPS reveals ultrastructural changes of mitochondria in acute kidney injury

To demonstrate the application of MAPS in studying mitochondrial dysfunction during disease progression, we analyzed ultrastructural changes in acute kidney injury (AKI). AKI is a multifactorial and multiphasic renal disease characterized by sudden loss of kidney function, leading to the accumulation of metabolic waste and toxins, often resulting in complications and failure of other organs^110^. Mitochondrial dysfunction has increasingly been identified as both an initiator of and contributor to AKI pathogenesis, as well as a therapeutic target^98,99^.

We investigated the impact of AKI on the 3D ultrastructure of mitochondria in cortical proximal tubular segments (S1/S2). AKI was induced in mice through treatment with cisplatin, a widely used cancer chemotherapy drug with a major side effect of nephrotoxicity^111^. Kidneys were collected from untreated (Control) and cisplatin-treated (AKI) mice, then processed using the pan-ExM-t workflow to generate 3D images. **Figure 6A** and **6B** show the mitochondria in the expanded cortical proximal tubules from control and AKI mice, respectively (**Video S5 and S6**). Qualitative visual inspection revealed not only morphological changes of the brush border and general cell morphology (**Figure S11**), but also that a substantial portion of mitochondria in AKI samples displayed disruption of the lamellar cristae we frequently observed in healthy renal proximal tubules. We next tested whether MAPS would be able to quantitatively confirm these anecdotal observations. MAPS successfully identified mitochondrial subcompartments with and without lamellar cristae. The high imaging speed (∼5 minutes/VOV) combined with the automated analysis, allowed us to image and analyze a total of ∼50 VOVs across ∼ 32 proximal tubules from 3 AKI and 3 control mice. Quantitative analysis of the identified 14,065 mitochondria from control samples and 10,227 mitochondria from AKI samples revealed a significant decrease in the fraction of lamellar cristae in AKI mice (**Figure 6C**): the average volume fraction containing lamellar cristae per mitochondrion and VOV dropped from 26% in controls to 0.7% in AKI samples (**Figure 6D**). This result illustrates the importance of the analysis of the mitochondria ultrastructure as no significant changes are observed with respect to the average length, volume or diameter (**Figure 6E-G, S12A-C**). However, as indicated by the sphericity analysis, mitochondria in AKI samples were significantly rounder than in control samples (**Figure 6H**). Across all of these metrics, we observed a large VOV-to-VOV variability (**Figure 6C-H**), emphasizing the need to image large numbers of VOVs to obtain reliable quantitative results.

**Figure 6.**
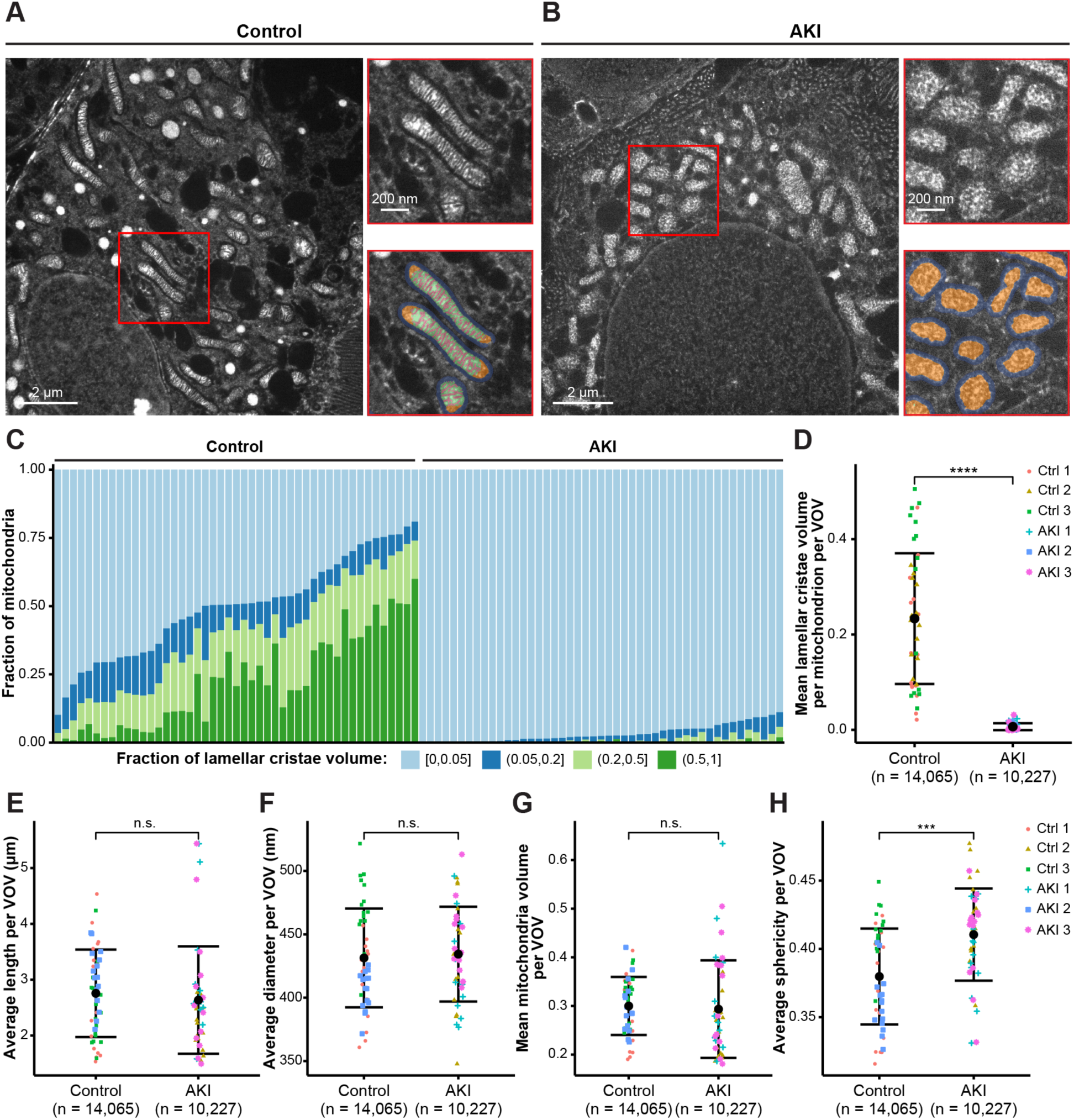
MAPS detects suborganellar morphological changes in mitochondria in acute kidney injury. **(A and B)** Representative images of expanded kidney tissue under control conditions (A) and after cisplatin treatment, which induces acute kidney injury (AKI) (B). Magnified views of the regions in the red boxes are shown without (top right) and with (bottom right) segmentation overlay next to the overview images. Segmentation highlights IMS (blue), matrix (green), cristae (pink), and non-lamellar ultrastructure areas (orange). Scale bars have been corrected for the expansion factor. **(C)** Distribution of the fraction of mitochondria with varying lamellar volumes (≤5%: light blue; 5-20%: blue; 20-50%: light green; >50%: green) in control and AKI samples. Each column represents one imaged volume. Control: *n* = 14,065 mitochondria from 47 imaged volumes, *N* = 32 proximal renal tubules from 6 independent experiments; AKI: *n* = 10,227 mitochondria from 50 imaged volumes, *N* = 33 proximal renal tubules from 6 independent experiments. Measurements were corrected for the expansion factor. **(D)** Mean lamellar cristae volume per mitochondrion per VOV. AKI samples show significantly less lamellar cristae compared to control samples. **(E-H)** Quantitative analysis of mitochondrial morphological changes in control and AKI samples. No significant differences were observed in average mitochondrial length per VOV (E), average diameter per VOV (F), or mean mitochondrial volume per VOV (G) between control and AKI samples. However, mitochondria in AKI samples displayed significantly reduced average sphericity per VOV (H) compared to control samples. Estimated means are shown in black, with error bars representing one standard deviation. ****, P < 0.0001 (Mann-Whitney Test); ***, P < 0.001; n.s., not significant. Control: *n* = 14,065 mitochondria from 47 imaged volumes, *N* = 32 proximal renal tubules from 3 mice in 6 independent experiments; AKI: *n* = 10,227 mitochondria from 50 imaged volumes, *N* = 33 proximal renal tubules from 3 mice 6 independent experiments. Measurements were corrected for the expansion factor.

## Discussion

Mitochondrial dysfunction is implicated in a large spectrum of diseases across nearly all fields of medicine. Yet, the mechanisms and morphological changes underlying this dysfunction and their role in disease progression remain in many cases poorly understood. Two challenges on the microscopy side have to be overcome to enable these currently lacking insights: one of contrast and one of throughput. With respect to the former, a microscopy method is needed that can resolve mitochondrial morphology in the context of the surrounding organelles and tissue as well as reveal molecules of interest therein. Regarding the latter, this imaging approach needs to be sufficiently accessible and fast to allow for studies in which tens to hundreds of samples can be imaged and quantitatively analyzed within weeks, not years. The presented automated workflow of MAPS that automatically segments mitochondrial subcompartments from 3D super-resolution pan-ExM images that can be rapidly obtained with a spinning disk confocal microscope offers a solution to both of these challenges.

The ease of sample preparation and the speed of 3D raw data acquisition in pan-ExM already surpassed established volume-EM - and in particular CLEM - approaches by large factors: a set of five to ten samples can be prepared in about a week without the need for highly specialized equipment. Imaging sample volumes consisting of ∼1,000 image frames can usually be acquired in less than 10 minutes on a standard spinning disk confocal microscope. However, having acquired the raw image data is only the first step towards the quantitative interpretation needed for systematic studies of organelle morphology across many samples. Additionally, a data analysis pipeline is required that can keep up with the high data acquisition speeds. MAPS’ DL-based approach provides such a pipeline. We chose U-Net as a core component since it is now an accepted reference architecture for semantic segmentation in biomedical imaging^112^.

Training such a model, however, represents a hurdle as high as those of sample preparation and data acquisition since the training process traditionally requires hundreds to thousands of hours of manual annotation to provide the necessary ground truth data. To overcome this second hurdle, MAPS takes advantage of pan-ExM’s unique correlative imaging strength which combines the cellular ultrastructure channel with channels showing specifically labeled targets that represent implicit, automatically co-registered annotations. This cuts down manual annotation from weeks or even months by one to three orders of magnitude to less than ten hours of effort.

Using the trained model, collecting and analyzing >10,000 mitochondria from ∼50 HeLa cells could then be achieved in less than 8 hours. Compared to pipelines that segment mitochondria using a combination of vEM or CLEM data and ML model^32,56–64^, the overall workflow of MAPS starting with sample preparation and ending with quantitative results is orders of magnitude higher throughput.

Related studies reported center-to-center distances between neighboring cristae of 70-185 nm and 71-227 nm on the basis of 10 and 15 mitochondria identified in TEM and STED 2D images, respectively^70^. While our results were in general agreement with these numbers when considering the different distance metric (center-to-center vs. surface-to-surface), ours are based on measurements obtained from 6,277 mitochondria in 3D data sets of 30 cells, reflecting a more than 100-fold increase in throughput.

Our larger numbers enabled us to reliably quantify even subtle changes at the organelle as well as the cristae level when cells were treated with oligomycin, a known ATP synthase inhibitor. MAPS could robustly detect both that mitochondria became shorter and rounder, as well as those cristae got sparser and less sheet-like - even if these changes were relatively small such as the inter-cristae distance increasing on average only by ∼6 nm (from ∼40 to ∼46 nm). By enabling this precise, large-scale 3D analysis, we believe that MAPS can provide insights into dynamic remodeling of mitochondria in response to other metabolic stressors such as hyperglycemia, deprivation, uncoupling, and diabetic conditions.

Proteomics resources, such as MitoCarta (now in its third iteration, MitoCarta3.0), MitoMiner, and MitoCoP, have been instrumental in identifying mitochondrial proteins in mammalian cells or tissues^15,16,113,114^. However, these methods for assigning proteins to submitochondrial localizations lack imaging data from studies on endogenous protein localization at the nanoscale. It is thus likely that some bona fide mitochondrial proteins are missing, and some annotations, particularly for less well-studied proteins, are incorrect. Meanwhile, atlases such as OpenCell (https://opencell.sf.czbiohub.org/) and The Human Protein Atlas (http://proteinatlas.org/) provide the subcellular maps of protein localization^115,116^ but lack the nanoscale resolution needed for precise insights into protein localization. These limitations highlight the need for advanced methods to directly visualize and quantify protein localization at the nanoscale. In this study, we demonstrate MAPS’ capability to do just that by localizing three well-known mitochondrial proteins, TOM20, MIC60, and COX4, within mitochondrial subcompartments. Seeing clear differences in their distributions, consistent with their known localization, this suggests that MAPS can enhance existing atlases by providing unbiased, high-throughput data on nanoscale mitochondrial protein localization.

Importantly, (mostly) removing the reliance on manual annotation reduces the influence of subjective judgement by the annotator which can introduce biases if not managed carefully. Combined with the substantially larger sample numbers that can now be processed, we believe this enables more reliable and robust quantification of structural features in heterogeneous samples. We demonstrated this here by imaging and quantifying 60 image stacks of mitochondria in proximal renal tubules taken from 3 cisplatin-induced mice experiencing acute kidney injury and 3 untreated mice, yielding data on more than 10,000 mitochondria for both the treated and untreated group of animals. Although previous studies reported the S3 segment being particularly sensitive to injury, the segment-specific alterations in metabolism in AKI remain poorly understood^103^. Our data on the S1/S2 segment found no significant change in the overall mitochondrial shape, but a significant decrease in the lamellar cristae volume ratio. These results underscore how advanced resolution and automated analysis can uncover subtle alterations in disease models. Importantly, our data also revealed a high level of heterogeneity across VOVs within the same sample group which highlights the value of unbiased imaging and analysis approaches that can be performed at high throughput.

Importantly, neither the sample preparation and data acquisition nor the quantitative analysis process of MAPS require any highly specialized equipment. This makes the presented approach easily portable to be scaled up and adopted by other research groups using the provided code, models, and guidelines. While tissue architecture varies significantly at the macroscopic level, cristae morphology seems to be remarkably conserved across different tissues. Based on our experience when transitioning from HeLa cells to kidney tissue, we expect that many applications of MAPS can be run with little (<10 hours) to no additional manual annotation and subsequent retraining on a large variety of tissues, such as the heart, liver, and brain, and cell types as long as similar pan-ExM sample preparation and labeling protocols are applied and images feature a comparable resolution, contrast and voxel size as our data.

While we have focused on mitochondria in this work, we anticipate that the presented approach can in the future be adapted to other cellular landmarks and organelles. Structures such as the nucleus, nuclear pore complexes, centrioles, and cilia, in particular, feature strong contrast and characteristic patterns in the pan imaging channel. For organelles for which this is not the case, e.g. the endoplasmic reticulum, alternative stainings that highlight lipids or other molecules present in bulk might create sufficient contrast.

Similarly, MAPS data analysis is expected to be compatible with other ExM protocols that are compatible with pan-stainings^36,39,40,44,117–119^. However, we expect that for most organelles’ expansion factors of 15 or more (or the combination with optical instruments offering resolutions substantially exceeding those of spinning disk confocal microscopes) will be needed to generate the necessary characteristic spatial features.

Independent of the described potential of MAPS as a general quantification tool, this study has demonstrated that the automatic segmentation of mitochondrial structures at scale is possible, which in itself is an important milestone. Like other reference data sets ^120^, our publicly available models, annotations, training data, imaging data, and segmentation results - to our knowledge the biggest dataset on mitochondria morphology to date - enable others to use our results as a resource.

## RESOURCE AVAILABILITY

### Lead contact

Further information and requests for resources and reagents should be directed to and will be fulfilled by the lead contacts, Jens Rittscher and Joerg Bewersdorf.

### Materials availability

This study did not generate new unique reagents.

### Data and code availability

- Data reported in this paper were deposited to BioImage Archive^121^ with accession number S-BIAD2200.
- The original MAPS code has been deposited at https://github.com/MAPS-ExM/MAPS, and the code for expansion homogeneity calculation is accessible at https://github.com/bewersdorflab/expansion-distortion-analysis. All code is publicly available.
- Any additional information required to reanalyze the data reported in this paper is available from the lead contact upon reasonable request.

## ACKNOWLEDGMENTS

We thank Ons M’Saad for valuable discussions on the human annotation of mitochondria in pan-ExM images. We thank Brett Phelan for his contribution in naming our methods and comments on the manuscript. We thank the support of the YALE ‘CINEMA’ Laboratory (Cellular Imaging using NEw Microscopy Approaches) for using the Andor Dragonfly High-Speed Confocal Microscope System. Y.T. and J.B. acknowledges support by the Wellcome Leap Foundation. A.S. is supported by the EPSRC Centre for Doctoral Training in Health Data Science (EP/S02428X/1). The computational aspects of this research were supported by the Wellcome Trust Core Award Grant Number 203141/Z/16/Z. J.R. is adjunct professor of the Ludwig Oxford Branch.

## AUTHOR CONTRIBUTIONS

Y.T., A.S., J.R. and J.B. designed the study. Y.T. optimized the pan-ExM protocol for mouse kidney tissue, prepared all samples, and performed image acquisition. A.S. performed the computational workflow for MAPS and quantitative analysis. Y.T., A.S., M.D., H.H., H.M. designed and performed manual mitochondrial annotations. Y.B. performed expansion homogeneity analysis. X.G., T.C., R.S., G.D. offered expertise in nephrology and provided fixed mouse kidney samples. J.R. and J.B. supervised the study. Y.T., A.S., J.R. and J.B. wrote the manuscript with input from all authors.

## DECLARATION OF INTERESTS

J.B. has licensed IP to Bruker Corp. and Hamamatsu Photonics. J.B. is a consultant for Bruker Corp. J.B. is a founder of panluminate, Inc. J.R. is a co-founder of Ground Truth Labs (Oxford, UK) and has received funding from Novo Nordisk.

## SUPPLEMENTAL INFORMATION

**Figure S1-S12**

**Video S1** Representative pan-ExM 3D image data overlapped with MAPS results of HeLa cells, related to Figure 2.

**Video S2-S3** Representative pan-ExM 3D image data overlapped with MAPS results of HeLa cells under different conditions: control (S2), and treatment with oligomycin (S3), related to Figure 4.

**Video S4** Representative pan-ExM 3D image data overlapped with MAPS results of proximal renal tubule, related to Figure 5.

**Video S5-S6** Representative pan-ExM 3D image data overlapped with MAPS results of proximal renal tubule under control conditions (S4) and acute kidney injury (AKI) (S5), related to Figure 6.

## STAR METHODS

### Key resources table

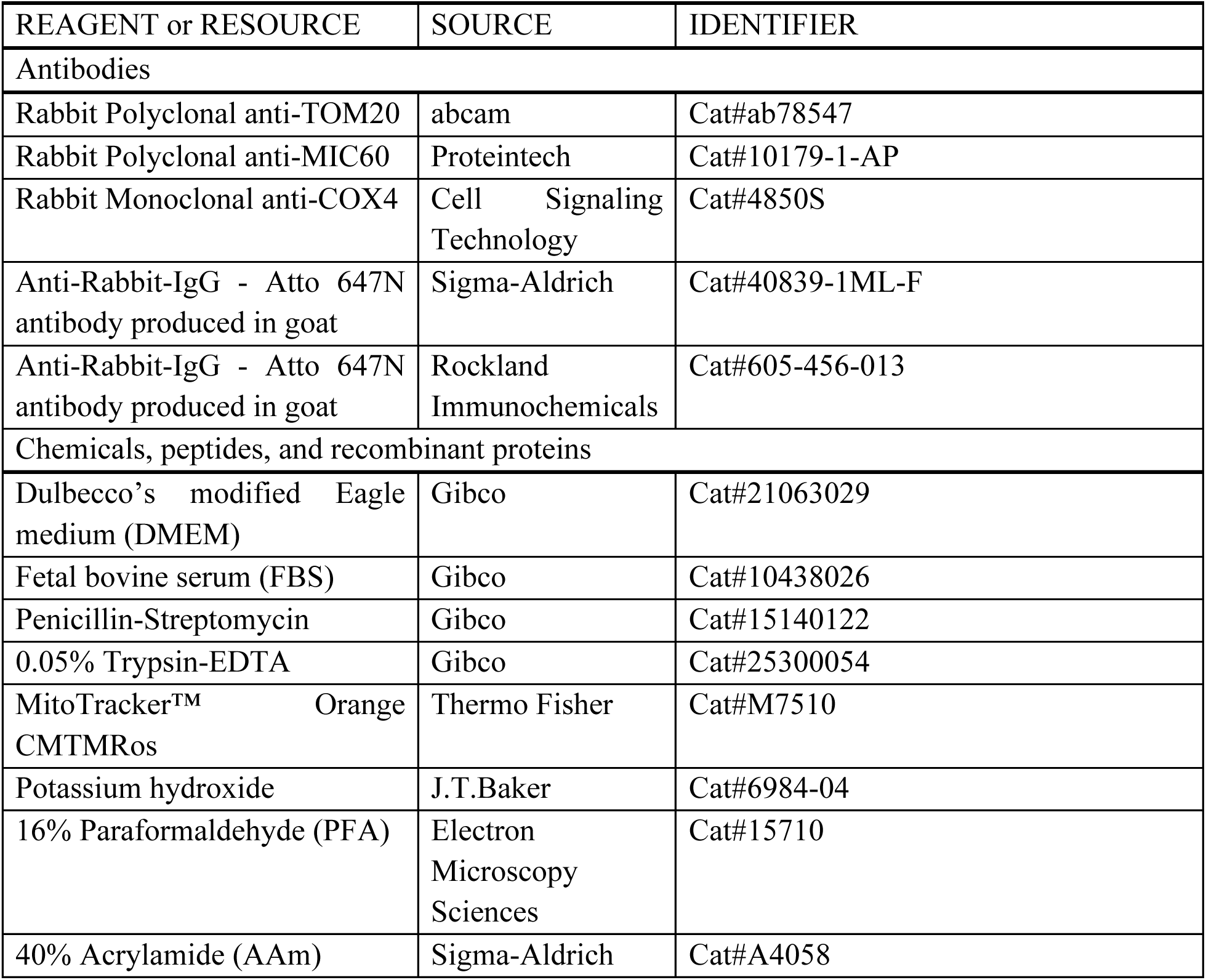
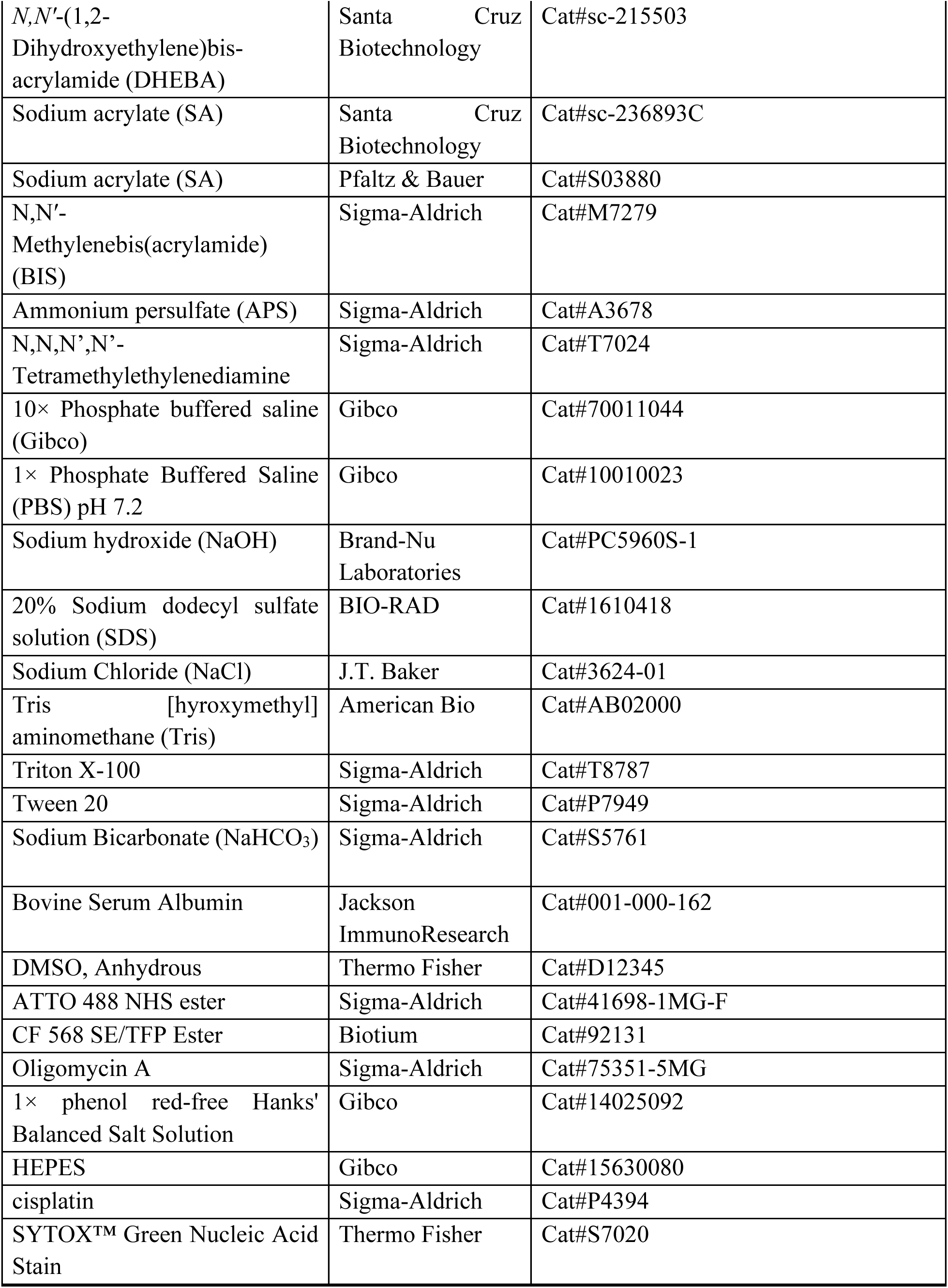
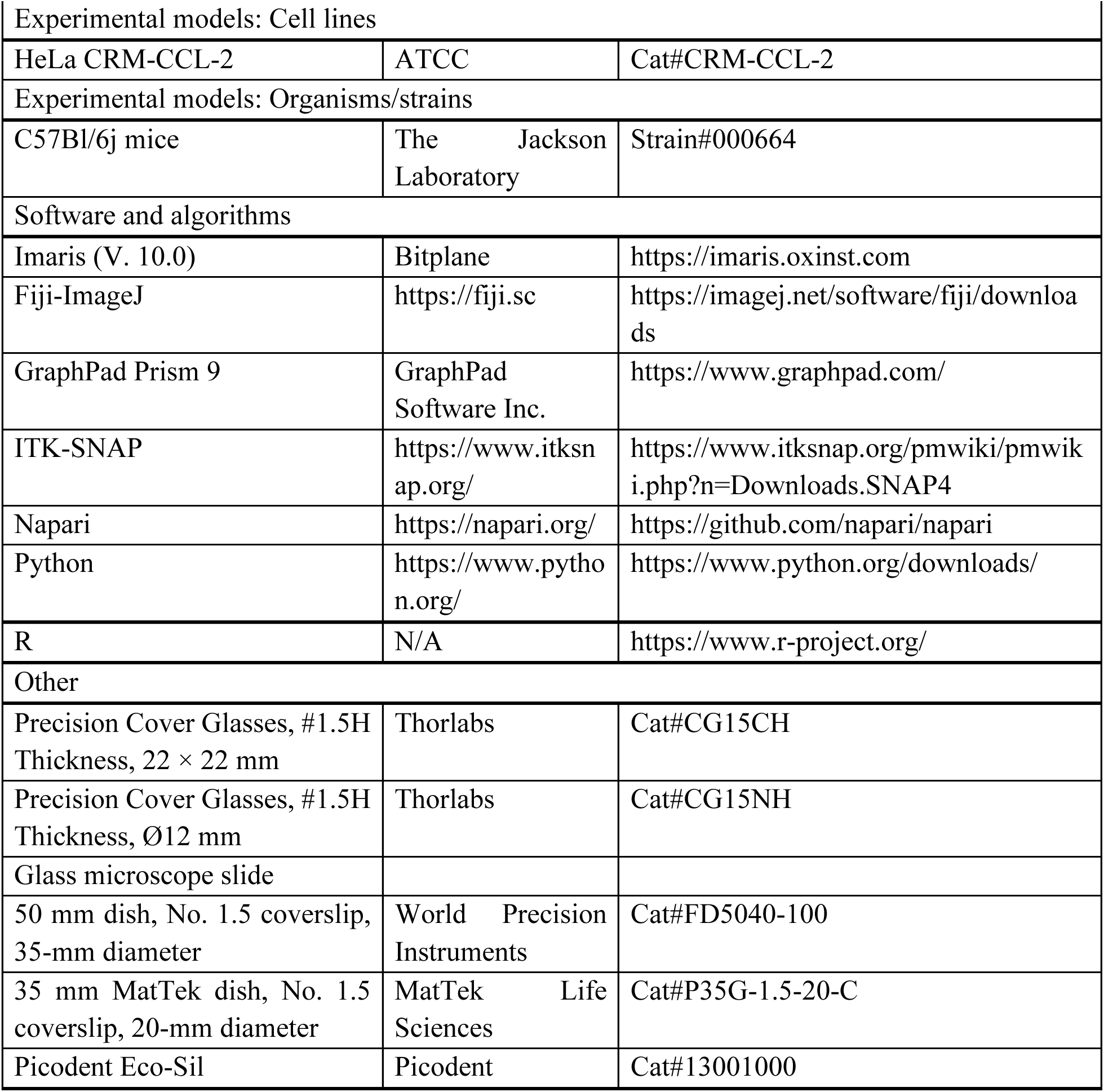

### EXPERIMENTAL MODEL AND STUDY PARTICIPANT DETAILS

#### Mouse lines

Mice were maintained at Veterans Affair Medical Center (VAMC, animal protocol number: RS0002), West Haven, CT. All experimental procedures were conducted according to the guidelines and regulations for animal care and use by the Institutional Animal Care and Use Committee of the VAMC and the authors complied with the ARRIVE guidelines. Mice were kept under a 12-h day/night cycle with food and water provided ad libitum. All experiments were repeated on at least two separate occasions. We used the ARRIVE1 reporting guidelines to report the animal studies.

#### Murine model of cisplatin-induced acute kidney injury (CP-AKI)

Adult male C57Bl/6j mice (∼4 months) were administered a single dose of CP (30 mg/kg) subcutaneously. Control mice were administered saline. Three days after CP treatment, blood samples were collected, and the mice were then sacrificed for kidney tissue collection. Plasma was prepared from the blood samples, and plasma creatinine levels were measured at the service of Yale animal physiology and phenotyping core to evaluate kidney function. The creatinine concentrations in the three control mice were 0.080, 0.064, and 0.068 mg/dL, respectively, and in the three CP-treated mice were 0.560, 0.384, 0.280 mg/dL, respectively.

### METHOD DETAILS

#### Cell culture

HeLa cells were grown at 37 °C, 5% (v/v) CO₂ in DMEM supplemented with 10% FBS and 1% penicillin-streptomycin at. Cells were passaged two to three times per week and used at passages below 20. Subculturing was performed using 1× PBS and 0.05% Trypsin-EDTA. Before cell plating, 12-mm round glass coverslips were cleaned in a sonic bath (Bronson) with 1M KOH for 15 minutes, followed by three rinses with MilliQ water. The coverslips were then sterilized with 100% ethanol, air-dried, and placed into 24-well plates. Next, cells were seeded onto the coverslips at ∼65,000 cells per well.

#### MitoTracker Orange staining

Before fixation, HeLa cells were incubated with 0.5 μM MitoTracker Orange for 30 minutes at 37 °C, 5% CO₂. The cells were then washed three times with culture medium and immediately fixed.

#### pan-ExM validated for expanding mammalian cells

pan-ExM was performed as previously reported^39^. Briefly, cells were fixed with 4% FA in 1× PBS for 1 h at room temperature and rinsed three times with 1× PBS. Samples were then post-fixed in 0.7% FA and 1% (w/v) AAm in 1× PBS for 6 h at 37 °C. After washing the samples three times with 1× PBS for 10 min each on a rocking platform, samples were embedded in the first expansion gel solution consisting of 19% (w/v) SA, 10% (w/v) AAm, 0.1% (w/v) DHEBA, 0.25% TEMED and 0.25% (w/v) APS in 1× PBS, and gelled for 1.5 hours at 37 °C in a humidified chamber. The samples were then transferred into a denaturation buffer containing 200 mM SDS, 200 mM NaCl, 50 mM Tris (pH 6.8) and incubated at 73 °C for 1 h. After washing out SDS and expanding the samples ∼4.5-fold in MilliQ water, gels were incubated in a re-embedding neutral gel solution (10% (w/v) AAm + 0.05% (w/v) DHEBA + 0.05% (v/v) TEMED + 0.05% (w/v) APS in 1× PBS) twice for 20 min each at room temperature. The residual gel solution was removed with Kimwipes, and the gels were sandwiched between a 22 × 22 mm coverslip and a glass microscope slide, followed by incubation at 37 °C for 1.5 h in a nitrogen-filled humidified chamber. The gels were rehydrated with 1× PBS and incubated in a second gel solution (19% (w/v) SA, 10% AAm (w/v), 0.1% (w/v) BIS, 0.05% (v/v) TEMED and 0.05% (w/v) APS) twice for 15 min each on ice. After removing the residual gel solution with Kimwipes, the gels were again sandwiched between a coverslip and a slide and incubated at 37 °C for 1.5 h in a nitrogen-filled humidified chamber. To dissolve DHEBA, the gels were incubated in 200 mM NaOH for 1 h at room temperature on a rocking platform, followed by three or four washes in 1× PBS. Next, gels were immunolabeled with primary antibodies diluted 1:250 in antibody dilution buffer (1% (w/v) BSA, 0.1% (v/v) Triton X-100 in PBS) for 24 h at 4 °C. Gels were washed in PBS-T (0.1% (v/v) Tween 20 in 1× PBS) three times for 20 min each at room temperature on a rocking platform. Next, samples were incubated with ATTO647N-conjugated anti-rabbit antibodies (diluted 1:250 in antibody dilution buffer) for 24 h at 4 °C. After washing with PBS-T three times for 20 min each, gels were incubated with 20 μg/mL NHS ester-ATTO488 or SE/TFP Ester-CF 568, and optionally SYTOX Green Nucleic Acid Stain (diluted 1:3000), dissolved in 100 mM NaHCO3 solution for 1.5 h, followed by three to five washes in PBS-T for 20 min each. Before imaging, the gels were expanded in MilliQ water to full expansion, mounted on glass-bottom dishes, and sealed with two-component silicone glue.

#### Oligomycin treatment

Cells were treated with 10 μM oligomycin A for 2 h, and control cells were treated with an equal volume of anhydrous DMSO. All cells were then fixed immediately after treatment and processed using pan-ExM.

#### Mice perfusion and kidney fixation

The mice were anesthetized by isoflurane and transcardially perfused with ice-cold 1× PBS, followed by 4% FA + 20% AAm in 1× PBS. Kidneys were then isolated and post-fixed overnight in 4% FA + 20% AAm in 1× PBS at 4 °C. After fixation, kidneys were washed twice in PBS, first for 1 h and then for 2 h at room temperature on a rocking platform, followed by an overnight wash at 4 °C on a rocking platform. Kidneys were subsequently stored in 1× PBS at 4 °C. The kidneys were sectioned at ∼50 μm using a vibrating microtome (Vibratome 1500, Harvard Apparatus). The sections were then washed four times for 15 min each in 1× PBS at room temperature and stored in 1× PBS at 4 °C for up to 1 month.

#### pan-ExM-t optimized for expanding kidney tissue sections

The pan-ExM-t protocol for expanding mouse kidney tissue sections was optimized based on the original protocol as previously reported^40^. Briefly, fixed kidney sections were incubated in inactivated first expansion gel solution (19% (w/v) SA, 10% (w/v) AAm, 0.1% (w/v) DHEBA in 1× PBS) for 30 min on ice. This was followed by incubation in an active first expansion gel solution (19% (w/v) SA, 10% (w/v) AAm, 0.1% (w/v) DHEBA, 0.075% TEMED and 0.0.075% (w/v) APS in 1× PBS) for 15 min on ice. The sections were then embedded in the active first gel solution and incubated for 2hr at 37 °C in a humidified chamber. The gels containing the sections were transferred to a denaturation buffer (200 mM SDS, 200 mM NaCl, 50 mM Tris, pH 6.8) and incubated at 73 °C for 6 h. After washing out SDS and expanding the samples ∼5-fold in MilliQ water, the gels were incubated in a re-embedding neutral gel solution (10% (w/v) AAm + 0.05% (w/v) DHEBA + 0.05% (v/v) TEMED + 0.05% (w/v) APS in 1× PBS) twice for 20 min each at room temperature. The residual gel solution was removed with Kimwipes, and the gels were sandwiched between a 22 × 22 mm coverslip and a glass microscope slide, followed by incubation at 37 °C for 1.5 h in a nitrogen-filled humidified chamber. After rehydration with PBS, the gels were incubated in a second gel solution (9% (w/v) SA, 10% AAm (w/v), 0.1% (w/v) BIS, 0.05% (v/v) TEMED and 0.05% (w/v) APS) twice for 15 min each on ice. The residual gel solution was removed with Kimwipes, and the gels were again sandwiched between a coverslip and a slide and incubated at 37 °C for 1.5 h in a nitrogen-filled humidified chamber. To dissolve DHEBA, gels were incubated in 200 mM NaOH for 1 h at room temperature on a rocking platform, followed by three or four washes in 1× PBS. Gels were then immunolabeled with primary antibodies (1:250 dilution in antibody dilution buffer: 1% (w/v) BSA, 0.1% (v/v) Triton X-100 in PBS) for 36 h at 4 °C. The gels were washed three times for 20 minutes each at room temperature and once overnight at 4 °C in PBS-T. The samples were then incubated with ATTO647N-conjugated anti-rabbit secondary antibodies (diluted 1:250 in antibody dilution buffer) for 36 hours at 4 °C. After washing three times for 20 min each at room temperature and once overnight at 4 °C on a rocking platform in PBS-T, the gels were incubated with 30 μg/mL NHS ester-ATTO488 or SE/TFP Ester-CF 568, and optionally SYTOX Green Nucleic Acid Stain (diluted 1:3000), dissolved in 100 mM NaHCO3 solution for 2.5 h, followed by three to five washes in PBS-T for 20 min each. Before imaging, the gels were expanded in MilliQ water to full expansion, mounted on glass-bottom dishes, and sealed with two-component silicone glue.

#### Image acquisition

Fluorescence images were performed using an Andor Dragonfly 600 High Speed Confocal Microscope Systems equipped with an inverted Nikon Eclipse Ti2 microscope (Nikon Instruments), an integrated laser engine (ILE), and Sona-6 sCMOS cameras. Images were acquired in spinning disk confocal mode (40-μm pinhole disk) using a CFI Plan Apo VC 60×, NA 1.2, water-immersion objective; an Apo LWD Lambda S 40×, NA 1.15, water-immersion objective; or a Super Fluor 20×, NA 0.75, air objective. Images were collected using 488 nm, 561 nm, and/or 637 nm excitation wavelengths, with image acquisition controlled through Fusion Control Software.

#### Image processing

3D renderings and maximum-intensity-projections were created using the napari python package. 2D images were created using the matplotlib python package.

#### Image Segmentation

The main objective of the proposed method is to minimize the required amount of human annotations by extracting all relevant information from the additional target-specific dyes (like MitoTracker for HeLa cells) or immunolabeling (MIC60 and COX4 for kidney samples).

Since the MitoTracker stains the mitochondria with high accuracy, traditional image-processing techniques are sufficient to derive target masks for the subsequent training following a similar approach to Stenman et al. (2021)^122^. Specifically, a median filter based on a 2-dimensional kernel disk with a size of 5 pixels allows to remove sufficient noise such that a subsequent morphological opening operation with a kernel size of 13 pixels generates mitochondria outline masks. The main objective of these masks is to fully contain all mitochondria as all volume outside of these masks is considered background in the subsequent model while any additional background within these masks is identified by the fine-tuned encoder. Moreover, a finer outline derived from the median filtered MitoTracker with a kernel size of 1 pixel is additionally added to allow the initial model to gather information about image features associated with the mitochondrial matrix. These masks correspond to the red and cyan segmentations found in **Figure 1B(i)**. The algorithm for the noisier immunolabeling is described below.

The base segmentation model follows a 3D U-Net architecture^66,67^ with residual connections^123^, group normalisation^124^ and weight standardisation^125^ which perform substantially better than batch normalization due to variations in the intensities of the pan staining. The number of internal channels of the encoder and decoder are 64, 128, 256 and 512. The UNet architecture was chosen due to its popularity to make the method accessible to researchers outside of the ML field and because it still performs on-par in most cases with other more complicated architecture^112^. We additionally perform extensive data augmentation in the training process in the form of intensity variation and geometric transformations like rotations and flipping. For the intensities, the maximum raw input is randomly clipped at different quantiles between 98.5% and 99.75% before normalizing to [0,1] in each training iteration to ensure robustness to variations in the mean intensity and account for different maximal intensities due to varying amounts the nucleus in the imaged volume of view.

Moreover, the intensities are randomly shifted by (x^α^+(1-(1-x)^β^))/2 with α and β between 1/5 and 5 to alternate the intensity histograms.

As a fully convolutional model, the model is not constrained to a specific input size but crops of 32×512×512 pixels are used in the training process. The initial pre-training based on the segmentation masks derived from the MitoTracker dye was conducted with twelve volumes of views with a size of 2048×2048 pixels and a depth varying between 71 and 380 slices with a mean of 210.

For the fine-tuning, the original encoder of the UNet is kept fix and a new decoder branch is trained based on 25 mitochondria crops were annotated that mainly cover one central mitochondrion and are of size 256×256 pixels with a depth between 20 and 35 slices of which 15 are used for training, five for validation and five for testing the final performance. The training terminates when no further improvement is seen on the separate validation set. These crops were annotated by five different annotators using the ITK-SNAP software (https://www.itksnap.org). The annotation process followed an annotation protocol that specified to first outline the IMS before overwriting the inner mitochondrion with the matrix label. Subsequently, the cristae label was added before a final step allowed to annotate ambiguous areas.

To compare the performance to other forms of pre-training for a smaller number of available training crops, the same architecture and training procedure is used but the first stage of the model is performed on the EM mitochondria dataset provided by Lucchi et al. (2012)^126^. Moreover, we also compare against a 2D version trained on ImageNet^127^. For every combination, three different models are trained based on different combinations of the reduced training and validation set and the mean and error bars are shown in **Figure S1A** demonstrating how the proposed approach reaches human-like performance with fewer required annotated crops. Human-like performance is defined as the annotator-to-annotator agreement in terms of the IoU metric. It was measured by three different annotators annotating eleven different mitochondria and resulted in an IoU of 0.34 due different interpretations of the boundaries of the mitochondria and the ambiguous areas.

3D UNets can be trained from sparse annotations^67^ in which only single slices are annotated instead of the full volume due to the high redundancy between adjacent slices. **Figure S1B** demonstrates that the effect of the initial training with the MitoTracker is especially beneficial in this case as the model is able to deduce the 3D feature distribution of the data from the MitoTracker masks instead of learning it from the annotated data. Moreover, **Figure S1B** shows that annotating only 40 slices distributed over 20 different mitochondria is sufficient to outperform a model trained from scratch based on over 200 slices over 20 mitochondria which substantially reduces the required annotated time.

For the final model the predictions of the initial outline model are used to disregard any background and the final segmentation masks are filled in by the additional fine-tuned decoder branch.

Moreover, for subsequent analysis an ensemble of six models trained on different training-validation splits for the HeLa cell model is employed with the final prediction given by a majority vote. Finally, possible segmentation mistakes are identified and refined^128^. To evaluate the final model, a human annotator reviewing and revising the model predictions for 30 mitochondria instances where necessary which resulted in an IoU value of 0.97. Moreover, in order to evaluate how well the model detects mitochondria we calculate the percentage of MitoTracker which is contained within our predictions compared to the overall amount of MitoTracker present in an image stack (**Figure S2**). After removing unspecific background labelling with a simple median filter, we find an agreement of 94% with a standard deviation of 2.6%.

In the case of the drug treated samples, a vector embedding for each treatment is learned which is then included in each layer of the de- and encoder and optimized during the training process. More specifically, for a drug, d, and corresponding embedding, z_d_, a two-layer MLP, f_θ_, is used to transform the intermediate features, X, of any convolutional block in the network by X _c_ = (1+f_θ_ ^scale^ (z_d_)_c_)*X_c_ + f_θ_ ^shift^ (z_d_)_c_ before the normalization is applied. This allows for different weightings of the internal features depending on the drug treatment but still enables the model to learn the features from all available data. The training procedure is repeated with additional segmentation masks derived from five drug-treated volume of views and 28 additional 256×256 pixel slices annotated from four crops for the ensemble training.

For the additional immunolabeling for the kidney tissue, it was not possible to derive the segmentation masks with the simple approach described above as the immunolabeling was not precise enough. Therefore the initial outline model is trained by casting the problem into a semi-supervised setting as shown in **Figure S9A**. More specifically, for each pixel i that is marked by the antibodies, a circular neighbor R_τ_(i) is defined and parameterized by some radius τ (which is set to twelve pixels for this work). The complement of the union of all neighbourhoods R(i) denoted by *U* serves as the background class. All pixels that are directly marked by the antibodies constitute the mitochondria class while all pixels in *U* that are not marked directly are considered unlabelled initially ignored in the loss function. This allows us to adopt a pseudo-labelling approach^129^. The model follows the same architecture described above and is trained to minimize the Dice loss and the Adam optimizer. After an initial warm-up phase of 500 gradient updates, pixels from *U* which are predicted with a high certainty are added to the corresponding class until the training stabilises. The threshold starts at 1 and is linearly interpolated down to 0.75 after 50.000 training steps. We observe that in general a low number of epochs like 6 is sufficient for good results. The resulting outline prediction is shown in **Figure S9B**. Subsequently, the outline predictions are filled with the ultrastructure-predictions from the previously described HeLa-model and a new model is trained to reproduce the combined predictions. Finally, the model is fine-tuned on sparsely annotated additional annotations. Because these annotations can be created efficiently by sparsely correcting the predictions of the previous predictions, we annotated three to five planes from 30 stacks of size 32×512×512 pixels resulting in 120 annotated slices requiring less than half a day of annotation effort. For the distinction between lamellar and non-lamellar ultrastructure, the fine-tuning process is repeated with annotations differentiating these areas **(Figure S9C)** in stacks from control and AKI samples. The required annotations can be completed even faster since the annotation structure is less fine-grained and in total 345 slices over 80 stacks were annotated in a few hours.

#### Parameter calculation

Individual mitochondria instances are identified based on separated inner membranes and then extended to include the IMS via a watershed algorithm^130^. The resulting mitochondria are skeletonized based on the classical axis skinning algorithm^131^ and post-processed by removing additional small side branches (shorter than 25 pixels) and dense artifacts areas. Moreover, the skeletons are linearly extrapolated to the boundary of each mitochondrion based on the direction of the last ten skeleton pixels. Additionally, side-branches for which the connecting line between end points are completely contained within the mitochondrion are removed. The length of the mitochondria is measured based on the resulting skeletons and the average diameter is calculated as twice the average distance from each pixel on the outer-membrane to the closest point on the skeleton.

#### Antibody identification

In order to investigate the localization of anti-TOM20, anti-MIC60 and anti-COX4, initial binding sites are identified by thresholding the intensities adaptively to match the mean binding number per mitochondria voxel between the antibodies. Subsequently, the center of each pixel cluster is identified as the location of the binding and clusters exceeding the 99% percentile size are split iteratively using k-Means. In order to measure the distance from the IBM, we calculate a signed distance map as shown in **Figure S5** and record the distance for every identified binding site. Moreover, we calculate the distance to the closest cristae junction for each antibody.

#### Expansion factor calculation

Nuclei in HeLa cells or kidney tissue sections from non-expanded and expanded using pan-ExM or pan-ExM-t were labeled with SYTOX Green Nucleic Acid Stain. Images were acquired using an Andor Dragonfly High Speed Confocal Microscope Systems with a 40×, NA 1.15, water-immersion objective or 20×, NA 0.75, air objective. Average nuclear cross-sectional areas were quantified using FIJI-ImageJ software. To calculate the expansion factor, the average nuclear cross-sectional area in the expanded samples was divided by the average cross-sectional area of non-expanded samples. The square root of this ratio was used as an estimate of the linear expansion factor. For mouse kidney sections, measurements were taken from the same nuclei before and after expansion. Results are presented in **Figure S3A** and **Figure S7B**.

#### Expansion homogeneity calculation

The spatial sample distortion was determined as previously reported^34^. In brief, the maximum projection of post-expansion and pre-expansion image stacks of nuclei were registered using open-source software Elastix. The post-expansion images were Gaussian blurred and registered to the corresponding pre-expansion images by similarity transformation. The similarity registered post-expansion images were then registered again to the pre-expansion images by B-spline transformation. The vector deformation field was extracted from the B-spline registration and applied to the binary outline of the nuclei. The distances between every two points on the outline before and after the deformation were calculated, and the error was measured by the absolute difference of the two distances. We measured the error for every combination of points across 6 μm on the outline of each nucleus. 10 nuclei from 6 independent experiments were analyzed.

### QUANTIFICATION AND STATISTICAL ANALYSIS

All statistical analysis was performed using R. Details of our statistical analysis can be found in the figure legends.

