## Supplementary Figures for "Scalable automated segmentation quantifies mitochondrial proteins and morphology at the nanoscale"

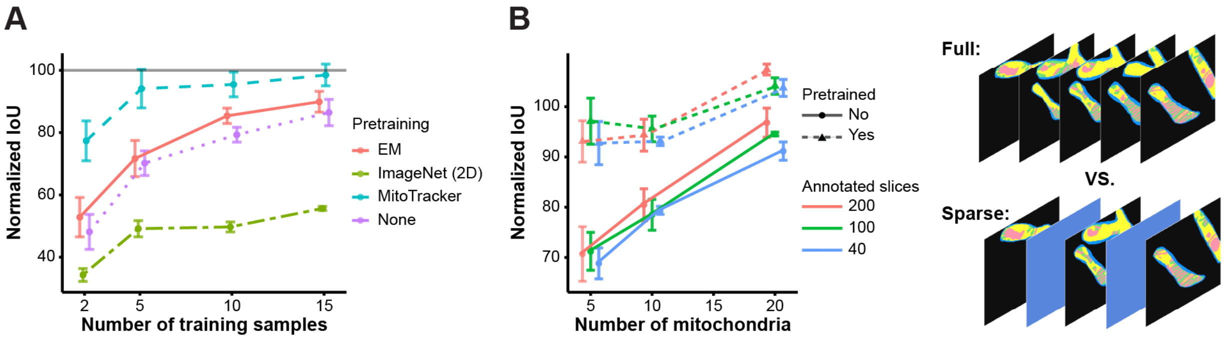


**Figure S1. Impact of pre-training with specific labeling and data sparsity on annotator agreement, related to Figure 1.**

**(A)** The percentage of annotator-to-annotator IoU agreement compared across four types of pretraining data: EM, ImageNet (2D), MitoTracker, and None (no pre-training). Error bars represent standard deviation across multiple samples. An annotator-to-annotator agreement of 100% corresponds to an IoU of 0.34 which was estimated by comparing annotations by three different annotators for eleven mitochondria. The MitoTracker pretraining shows the highest IoU with fewer training samples, followed by EM, no pre-training, and ImageNet (2D), respectively.

**(B)** The percentage of annotator-to-annotator IoU agreement compared across the number of mitochondria annotated across varying annotated slices (200, 100, and 40 slices) demonstrating that is far more effective to annotate larger variety of mitochondria sparsely than fewer mitochondria dense. Moreover, the MitoTracker pre-training improves the performance significantly over all combinations.

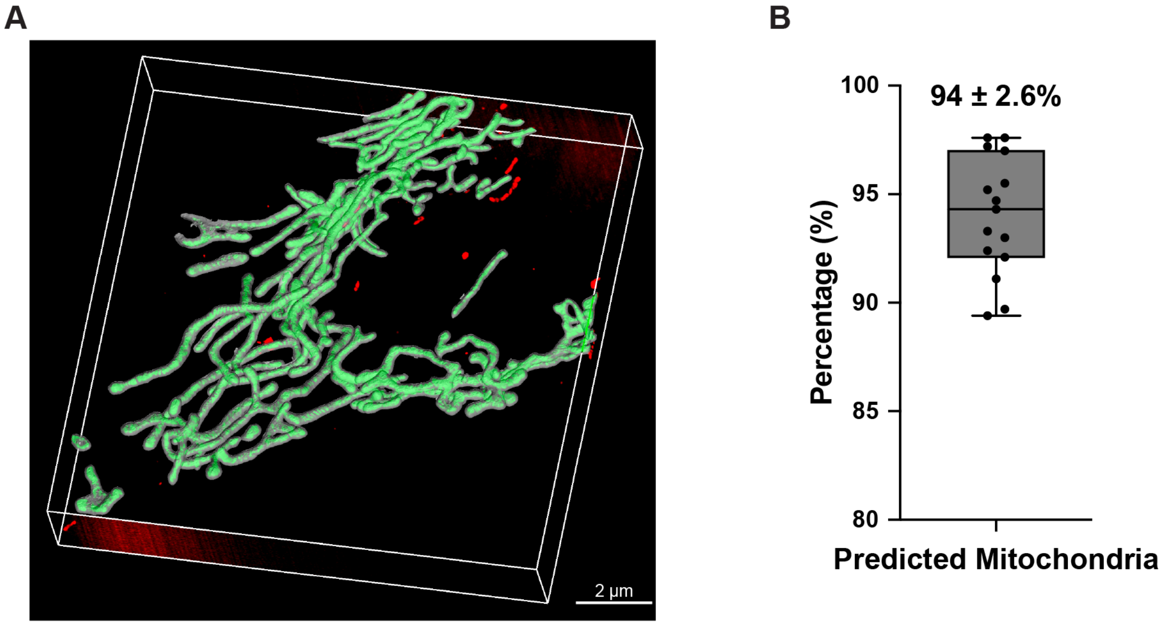


**Figure S2. Identification of mitochondrial volume using MAPS, related to Figure 2.**

**(A)** A 3D volumetric view of reconstructed mitochondria from an expanded HeLa cell illustrates the successful segmentation of mitochondrial structures (green) from the surrounding cytosol using MAPS. Regions not classified as mitochondria are shown in red.

**(B)** MAPS successfully identified mitochondrial regions with an average prediction accuracy of 94% ± 2.6%. Each dot represents one of 15 individual image volumes, with the box plot summarizing the distribution of prediction percentages.

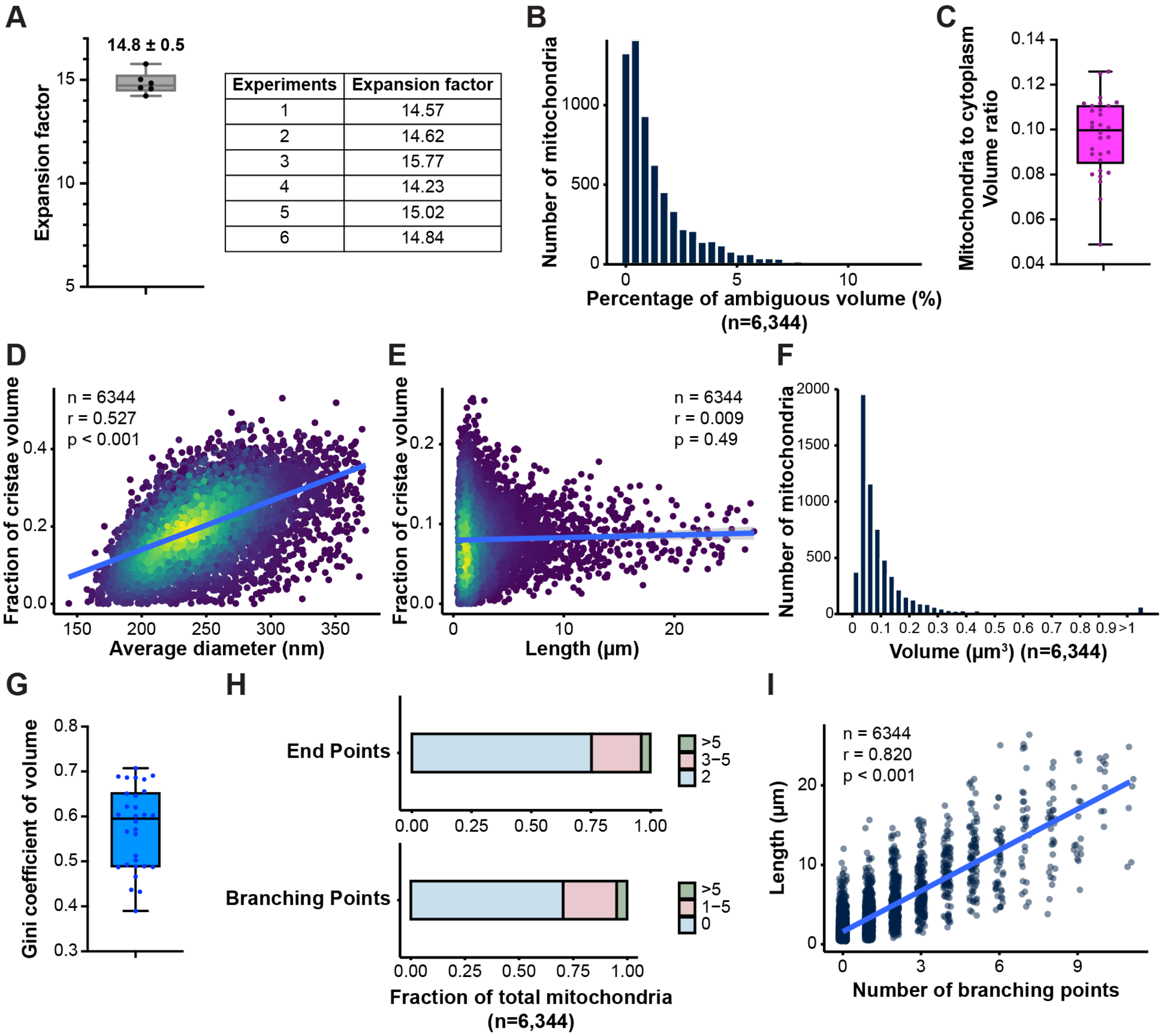


| Experiments | Expansion factor |
|---|---|
| 1 | 14.57 |
| 2 | 14.62 |
| 3 | 15.77 |
| 4 | 14.23 |
| 5 | 15.02 |
| 6 | 14.84 |

**Figure S3. Additional quantitative analysis of mitochondrial structure and ultrastructure in HeLa cells, related to Figure 2.**

**(A)** Estimated expansion factors achieved with pan-ExM were determined from averaged nuclei cross-section measurements comparing pre-expanded and post-expanded HeLa cell images (see **STAR Methods**). The mean expansion factor was calculated as 14.8 ± 0.5 (*n* = 140 nuclei from 6 independent experiments).

**(B)** Distribution of the percentage of ambiguous volume in mitochondria (*n* = 6,344 mitochondria, *N* = 30 cells from 4 independent experiments), showing 98% of mitochondria contained less than 5% ambiguous volume.

**(C)** Box plot showing the mitochondria to cytoplasm volume ratio in 3D pan-ExM images. The median and interquartile range are shown with whiskers extending to the minimum and maximum values, illustrating a mean ratio of 9% (*N* = 30 cells from 4 independent experiments).

**(D and E)** Scatter plot illustrating the relationship between mitochondrial cristae volume fraction and average mitochondrial diameter or length. A positive correlation is evident between mitochondrial diameter and cristae volume fraction, indicating that thicker mitochondria generally contain more cristae. In contrast, no clear correlation is observed between mitochondrial length and cristae volume fraction. n: number of analyzed mitochondria; r: Pearson's correlation coefficient; p: P-value.

**(F)** Distribution of mitochondrial volumes (*n* = 6,344 mitochondria, *N* = 30 cells from 4 independent experiments), showing 80% of mitochondria are 0.03-0.25 $\mu m^3$ in volume.

**(G)** Box plot of the Gini coefficient for mitochondrial volume, providing a measure of inequality in mitochondria distribution with a mean of 0.53 (*N* = 30 cells from 4 independent experiments).
**(H)** Fraction of mitochondria classified by the number of end points (top) and branching points (bottom). Most mitochondria show two end points and no branching points.
**(I)** Scatter plot showing the relationship between mitochondrial length and the number of branching points. A positive trend is observed with each branching point increasing the length on average by 1.72. n: number of analyzed mitochondria; r: Pearson's correlation coefficient; p: P-value.

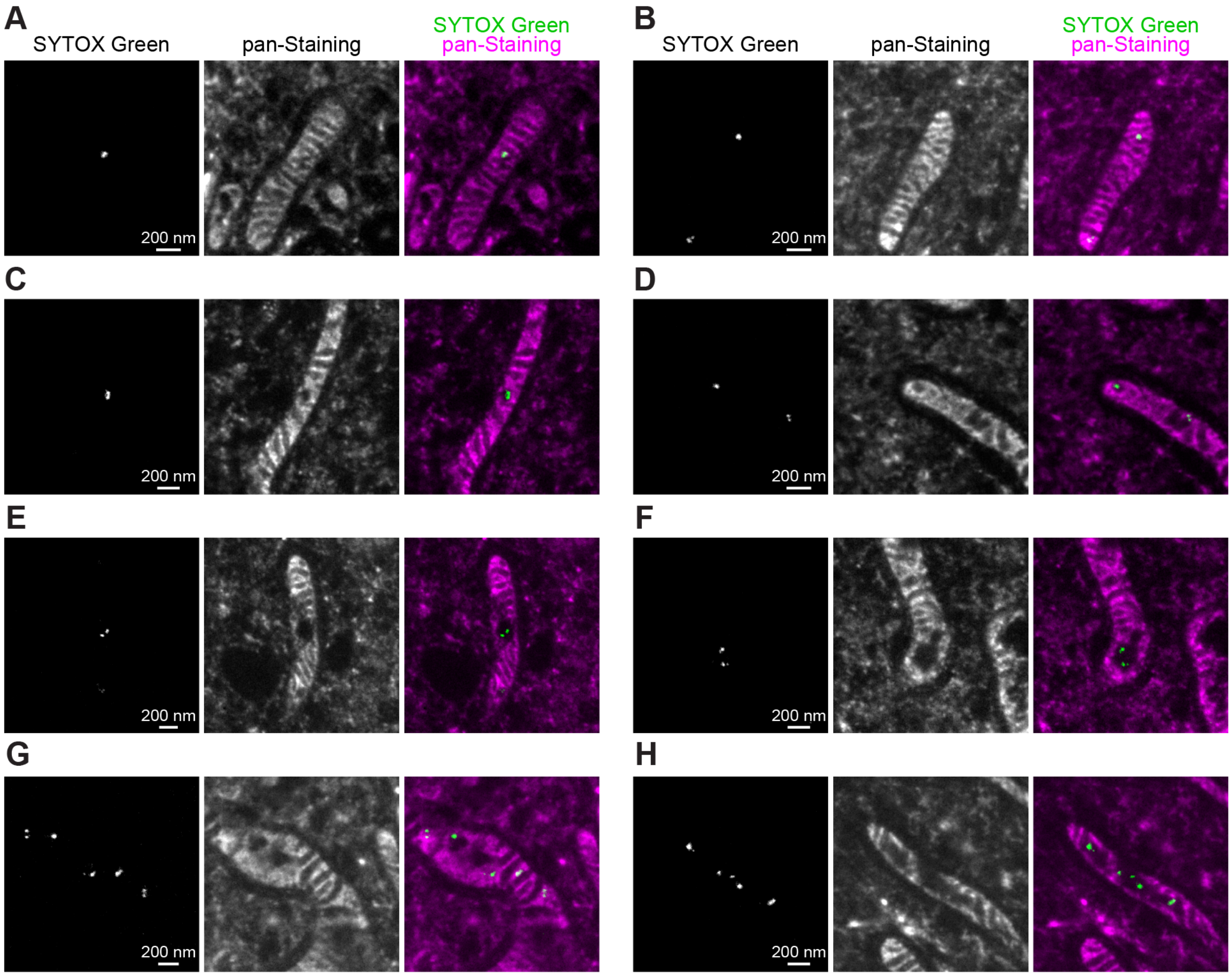


**Figure S4. Distribution of mitochondrial DNA within mitochondrial ultrastructure in expanded HeLa cells, related to Figure 2.**

Representative pan-ExM images showing mitochondrial DNA (mtDNA) labeled with SYTOX Green (left column), pan-staining (middle column), and their merged overlays (right column).

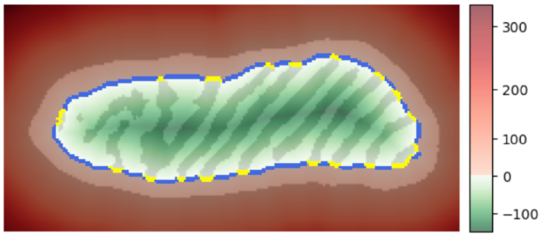


**Figure S5. Identification of IBM and cristae junctions, related to Figure 3.**
Heatmap representation of a segmented mitochondrion. The background color gradient represents the signed distance from the inner membrane in nm, as shown in the color bar. A distinct contour (blue) represents the inner boundary membrane (IBM), while yellow markers highlight cristae junctions.

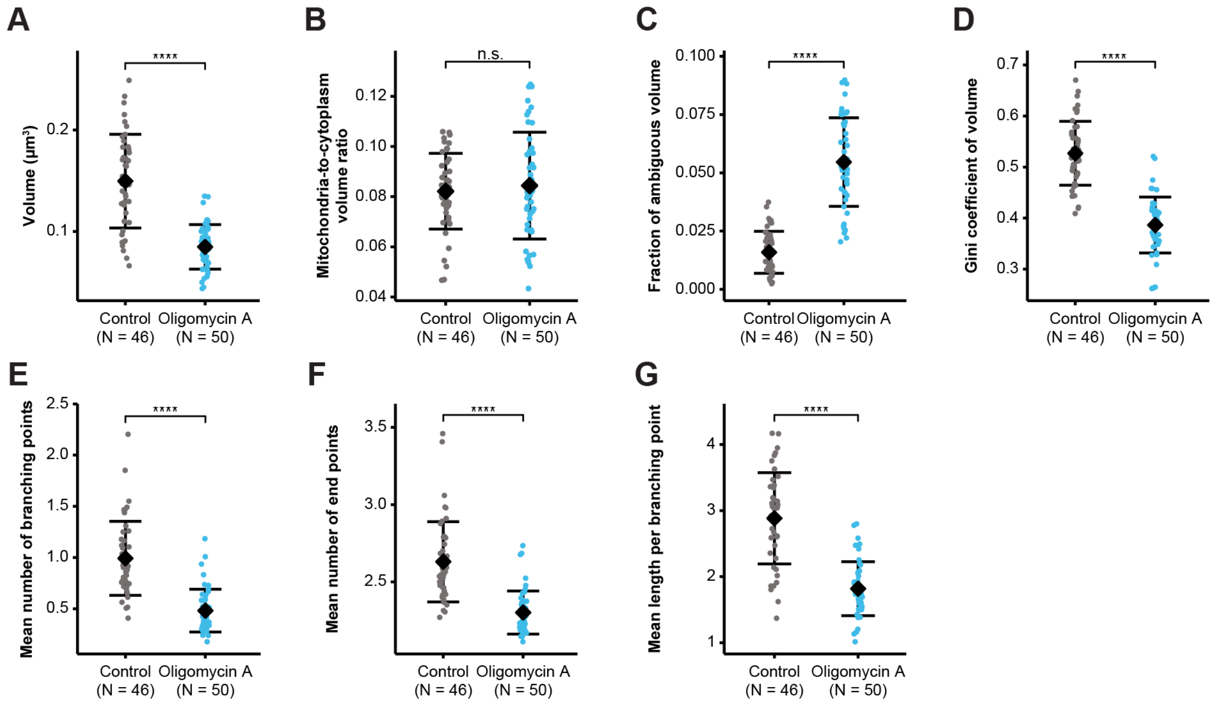


**Figure S6. Additional organellar and suborganellar morphological changes in mitochondria in oligomycin A-treated HeLa cells, related to Figure 4.**

Quantitative analysis of mitochondrial morphological changes under the different conditions (control: $n$ = 8,067 mitochondria, $N$ = 46 cells; oligomycin A: $n$ = 15,329 mitochondria, $N$ = 50 cells. All derived from 6 independent experiments). Estimated means are shown in black, with error bars representing one standard deviation. ****, $P < 0.0001$ (Mann-Whitney Test); n.s., not significant. Oligomycin A treatment results in significant reductions in total volume (A), the mean number of mitochondrial branching and end points (E and F), and mean length per branching point (G). Additionally, it leads to increases in ambiguous volume (C) and heightened equality in volume distribution (D), while the mitochondria-to-cytoplasm volume ratio (B) remains statistically indifferentiable.

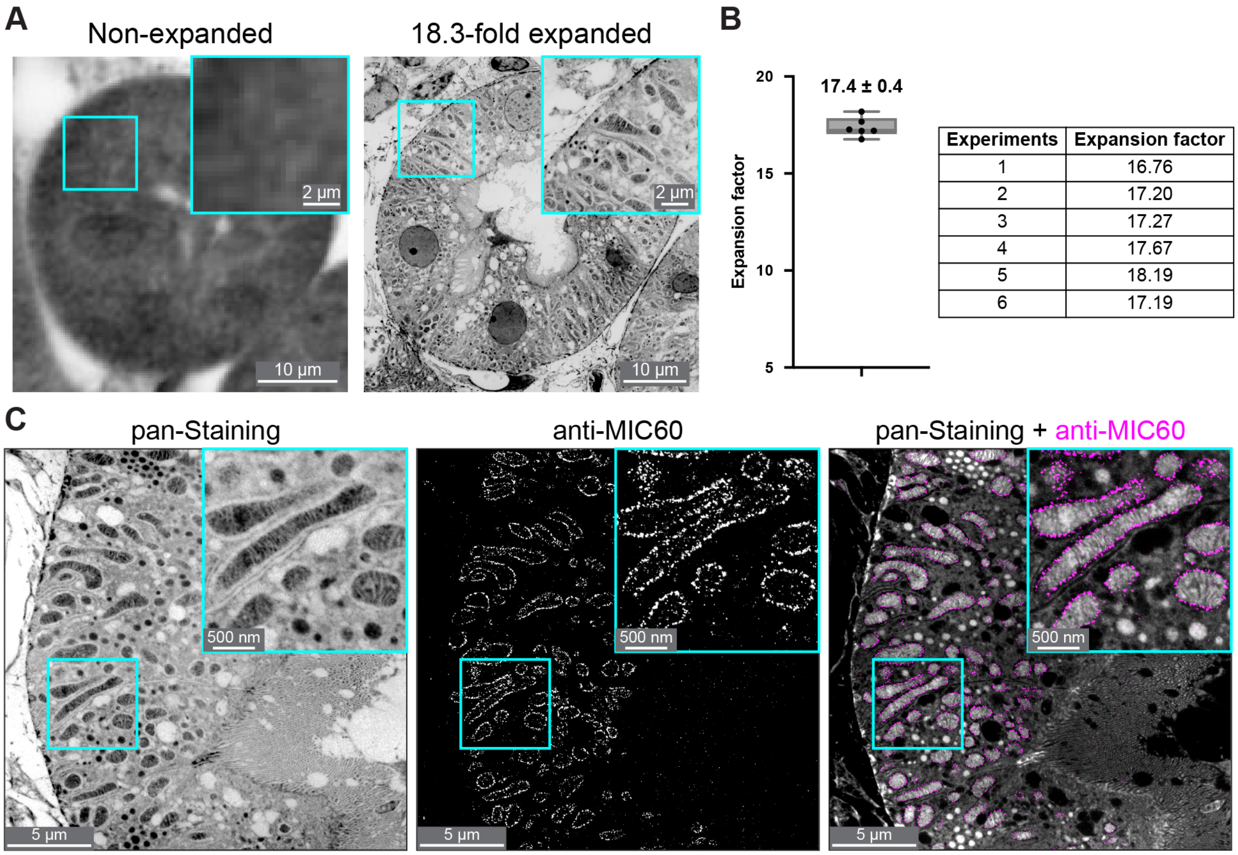


| Experiments | Expansion factor |
|---|---|
| 1 | 16.76 |
| 2 | 17.20 |
| 3 | 17.27 |
| 4 | 17.67 |
| 5 | 18.19 |
| 6 | 17.19 |

**Figure S7. pan-ExM expands mouse kidney sections and is capable of immunolabeling, related to Figure 5.**

**(A)** Representative cross-sectional images of the renal tubule before and after 18.3-fold expansion.

**(B)** Estimated expansion factors achieved with pan-ExM-t were determined by comparing the cross-sectional areas of the same nuclei in mouse kidney sections before and after expansion (see **STAR Methods**). The mean expansion factor was calculated as 17.4 ± 0.4 ($n$ = 291 nuclei from 6 independent experiments).

**(C)** Representative images of immunolabeling of mitochondria in an expanded mouse renal tubule. Left: pan-Staining showing overall mitochondrial structures, displayed with a white-to-black color table. Middle: Anti-MIC60 immunostaining, specifically labeling a component of the MICOS complex. Right: Merged image of pan-staining (gray) and anti-MIC60 (magenta).

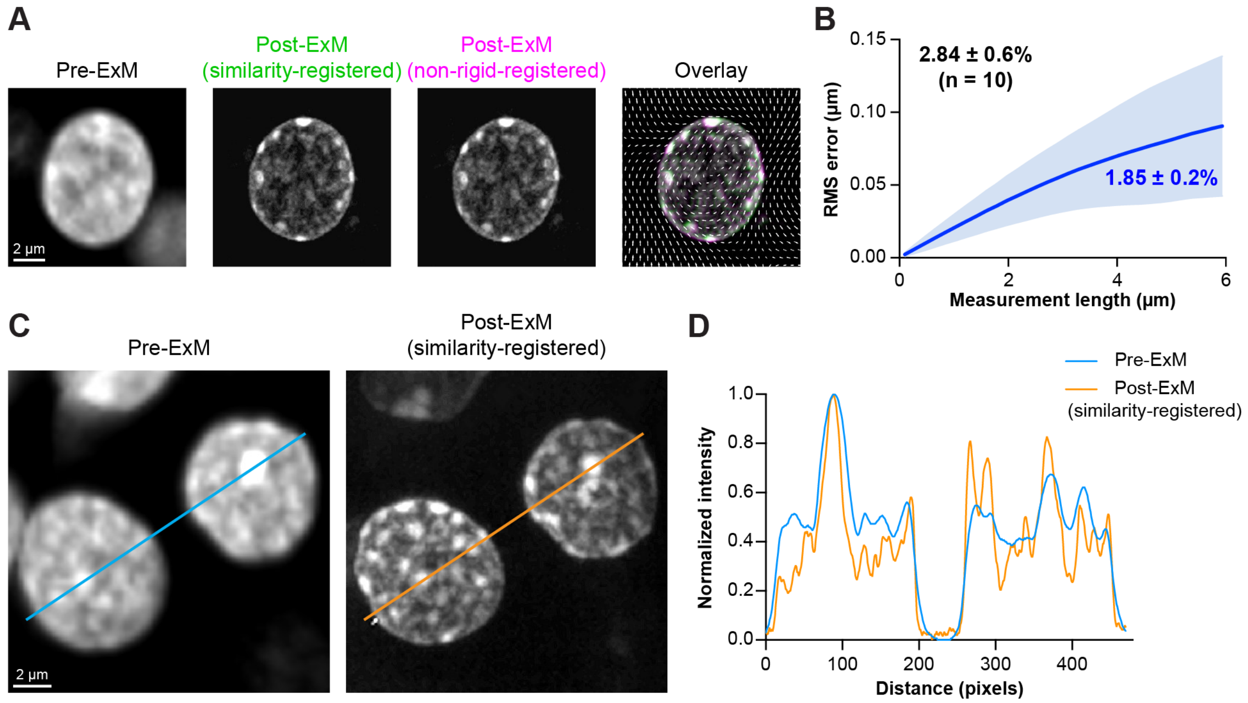


**Figure S8. RMS error homogeneity calculation in expanded mouse kidney sections using pan-ExM, related to Figure 5.**
**(A)** Assessment of sample deformation in nuclei before (Pre-ExM) and after expansion (Post-ExM). The leftmost column shows a pre-expanded nucleus. The middle columns display post-expanded nuclei after similarity-based registration (green) and non-rigid registration (magenta). The rightmost column overlays these registered images, with white arrows indicating the distortion vector field.
**(B)** Root-mean square (RMS) error quantification as a function of distance. The dark blue line represents the mean, and the light blue error bars correspond to the standard deviation. For structures 6 µm apart, the RMS error measured for the nucleus in (A) was 1.85 ± 0.2% of the measurement length. Across 10 nuclei (*N* = 10 proximal renal tubules from 3 mice in 6 independent experiments), the RMS error was 2.84 ± 0.6%. See STAR Methods for details on RMS error quantification.
**(C)** Comparison of intensity profiles between pre- and post-expanded nuclei after similarity-based registration, with blue and yellow lines marking line scan positions.
**(D)** Corresponding normalized line-scan plots of the lines drawn in (C), showing overall alignment in intensity distribution of pre- and post-expanded nuclei.

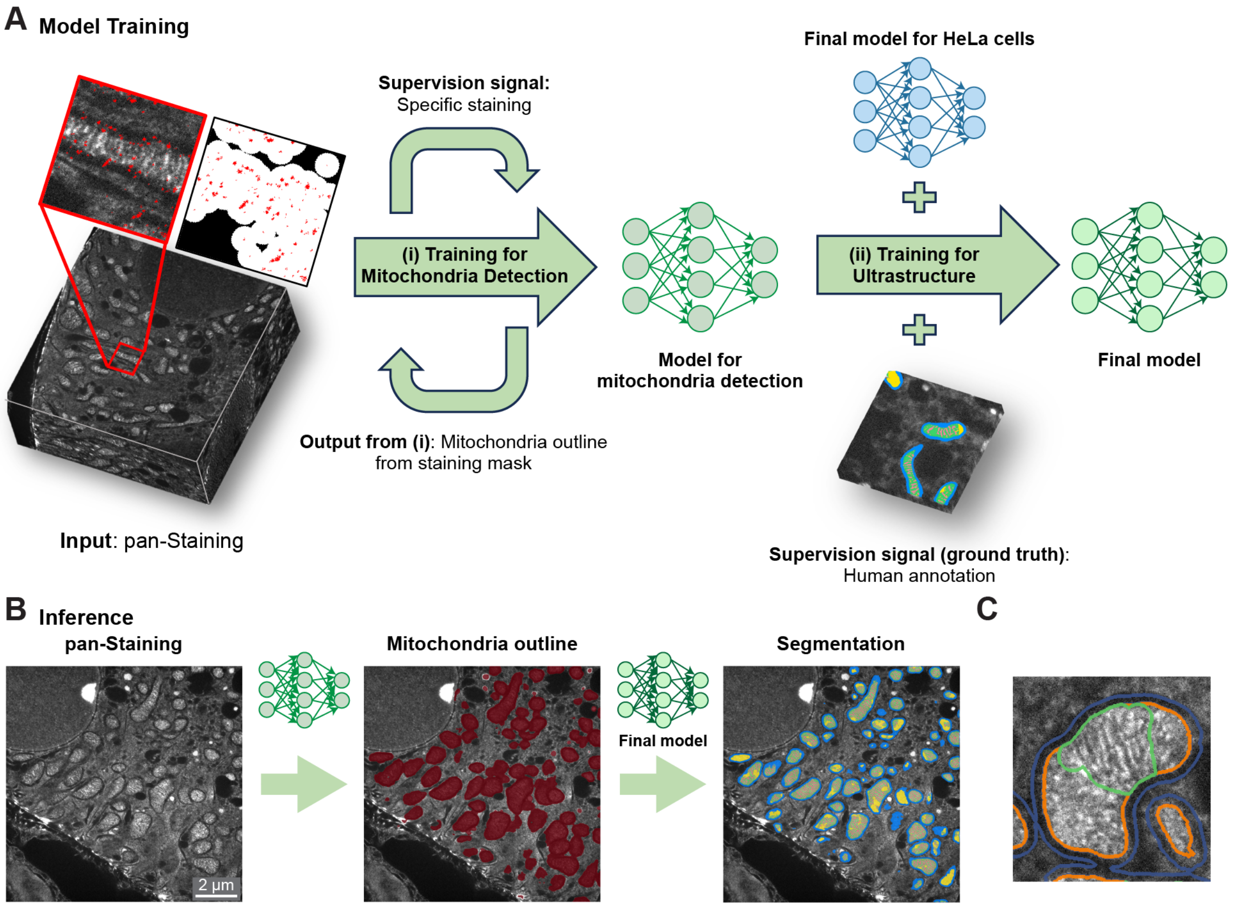


**Figure S9. MAPS workflow for mouse kidney sections, related to Figure 5.**
**(A)** Computational pipeline for training a model to segment mitochondrial subcompartments in kidney samples.
**(B)** The final model resulting from the training facilitates automated large-scale segmentation and quantitative analysis of mitochondria based solely on the pan-staining image. Scale bars have been corrected for the expansion factor.
**(C)** Example annotation of the distinction between lamellar cristae (green) and non-lamellar ultrastructure areas (orange).

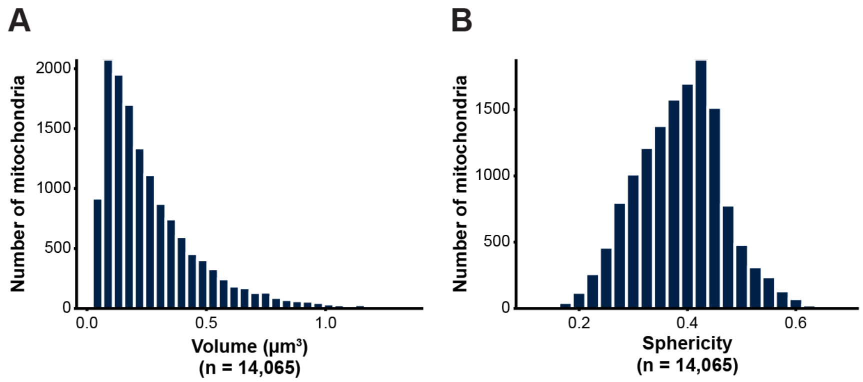


**Figure S10. Additional quantitative analysis of 3D mitochondrial morphology in mouse kidney tissue, related to Figure 5.**

Distribution of mitochondrial volumes (A) and sphericity (B), showing a volume of 0.29 ± 0.27 $\mu m^3$ (mean ± SD) and a sphericity of 0.39 ± 0.08 (mean ± SD). A total of *n* = 14,065 mitochondria were analyzed from 47 imaged volumes; *N* = 32 proximal renal tubules sampled from 6 independent experiments. Measurements were corrected for the expansion factor.

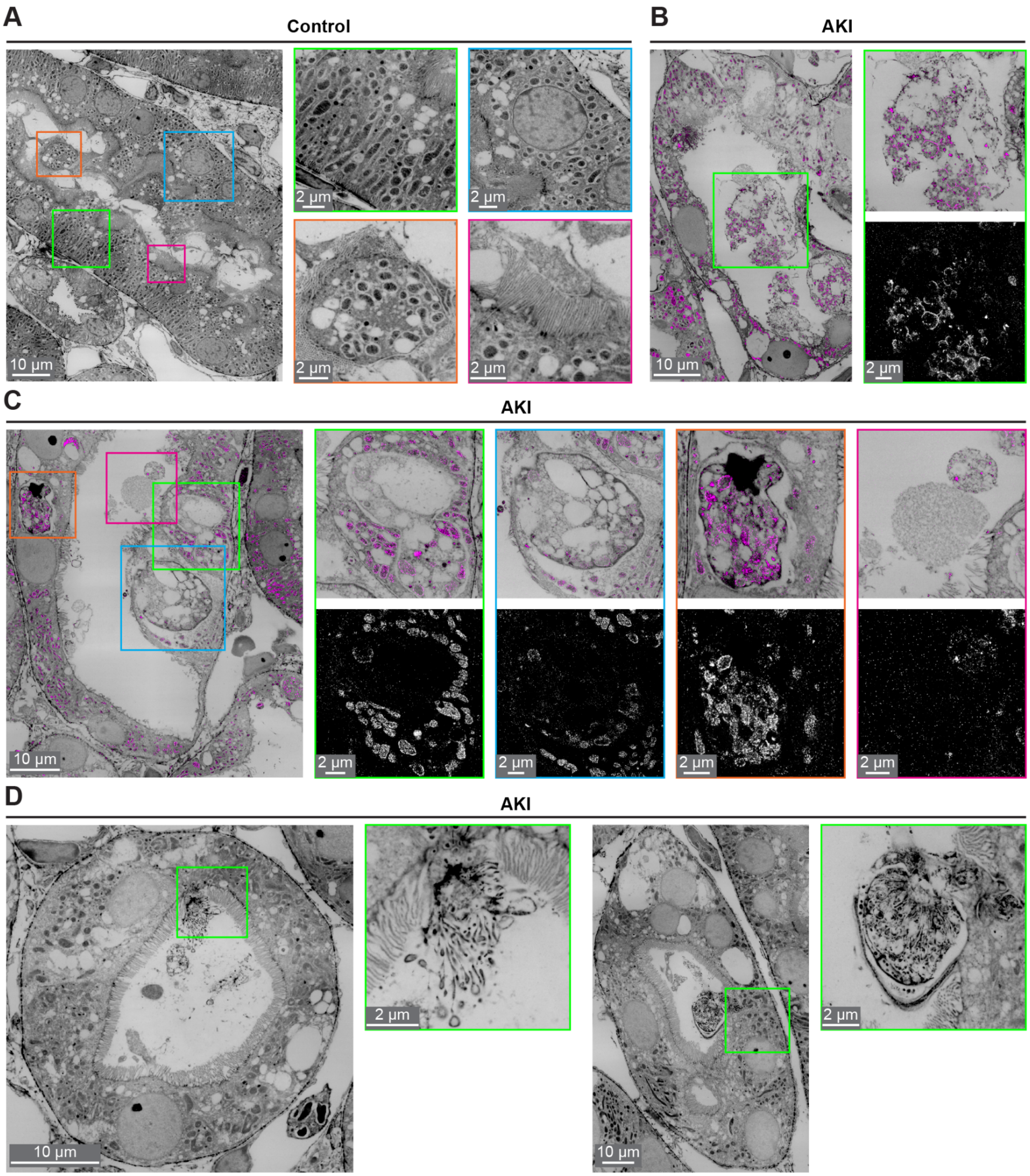


**Figure S11. Morphological changes in acute kidney injury, related to Figure 6.**
**(A)** Representative images of morphology of mitochondria and brush border in expanded kidney tissue under control conditions.
**(B-D)** Representative images of morphology changes of cell structure and the brush border in expanded kidney tissue after cisplatin treatment, which induces acute kidney injury (AKI). Proximal tubules in B and C were immunolabeled for the mitochondrial proteins MIC60 and COXIV in magenta.

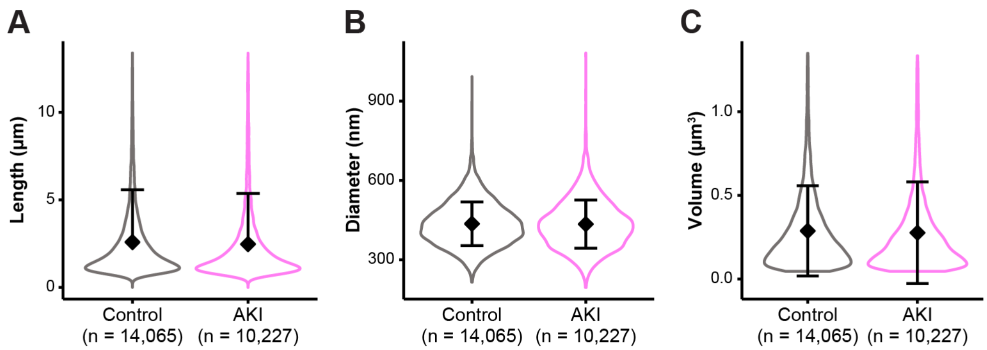


**Figure S12. Additional suborganellar morphological changes in mitochondria in acute kidney injury, related to Figure 6.**

Quantitative analysis of mitochondrial morphological changes in control and AKI samples. No significant differences were observed in average mitochondrial length per mitochondria (A), average diameter per mitochondria (B), or mean mitochondrial volume per mitochondria (C) between control and AKI samples. Control: *n* = 14,065 mitochondria from 47 imaged volumes, *N* = 32 proximal renal tubules from 3 mice in 6 independent experiments; AKI: *n* = 10,227 mitochondria from 50 imaged volumes, *N* = 33 proximal renal tubules from 3 mice 6 independent experiments. Measurements were corrected for the expansion factor.
